# Development of a Covalent Inhibitor of Gut Bacterial Bile Salt Hydrolases

**DOI:** 10.1101/640086

**Authors:** Arijit A. Adhikari, Tom C. Seegar, Scott B. Ficarro, Megan D. McCurry, Deepti Ramachandran, Lina Yao, Snehal N. Chaudhari, Sula Ndousse-Fetter, Alexander S. Banks, Jarrod A. Marto, Stephen C. Blacklow, A. Sloan Devlin

## Abstract

Bile salt hydrolase (BSH) enzymes are widely expressed by human gut bacteria and catalyze the gateway reaction leading to secondary bile acid formation. Bile acids regulate key metabolic and immune processes by binding to host receptors. There is an unmet need for a potent tool to inhibit BSHs across all gut bacteria in order to study the effects of bile acids on host physiology. Here, we report the development of a covalent pan-inhibitor of gut bacterial BSH. From a rationally designed candidate library, we identified a lead compound bearing an alpha-fluoromethyl ketone warhead that modifies BSH at the catalytic cysteine residue. Strikingly, this inhibitor abolished BSH activity in conventional mouse feces. Mice gavaged with a single dose of this compound displayed decreased BSH activity and decreased deconjugated bile acid levels in feces. Our studies demonstrate the potential of a covalent BSH inhibitor to modulate bile acid composition in vivo.

## Introduction

Bile acids, long relegated to the role of undistinguished detergents, have recently emerged as likely candidates for the molecular messengers that allow members of the human gut microbiome to modulate the physiology and behavior of their hosts.^1,2^ Investigating these potential roles in detail has been hampered by the lack of tools to regulate specific messages, and developing the appropriate tools has been hampered in turn by the complex biosynthesis of bile acids. Primary bile acids are produced in the liver from cholesterol and conjugated to taurine or glycine to produce primary conjugated bile acids (Fig. 1a). These molecules are stored in the gallbladder and released into the digestive tract where they aid in absorption of lipids and fat-soluble vitamins. Over 95% of bile acids are reabsorbed in the ileum and transported to the liver. The remaining ~5% pass into the colon, where the majority of gut bacteria reside. Gut bacteria then enzymatically modify these primary bile acids, producing a group of molecules called secondary bile acids (Fig. 1a). Roughly 50 secondary bile acids have been detected in human feces, and their concentrations can reach low millimolar levels.^3,4^

**Figure 1.**
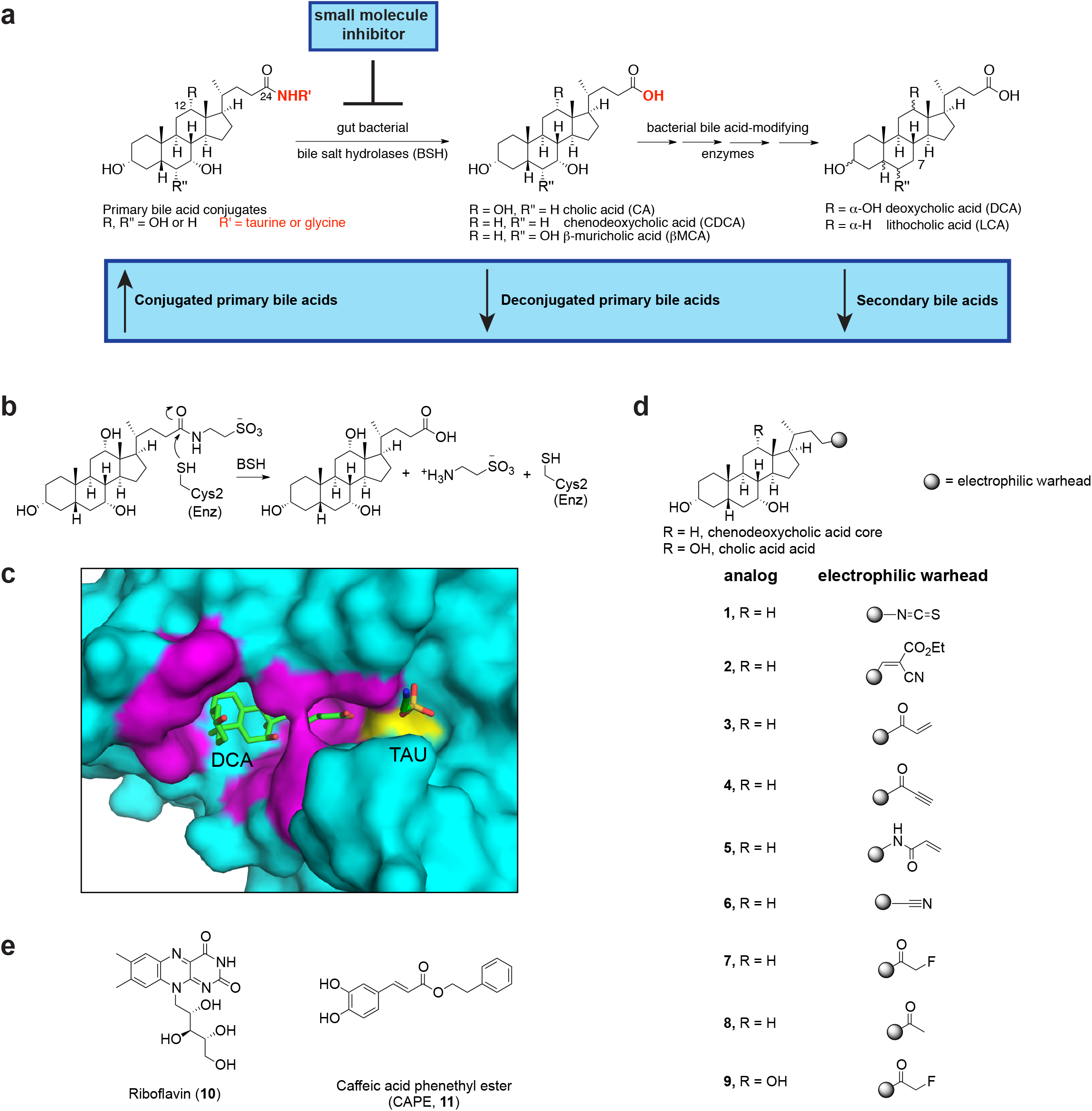
Rational design of small molecule inhibitors of gut bacterial bile salt hydrolase (BSH) enzymes. (**a**) BSHs are the gateway enzymes in the conversion of primary (host-produced) to secondary (bacterially produced) bile acids. Inhibition of BSHs should result in a decrease in deconjugated primary and secondary bile acids. (**b**) Mechanism of enzmyatic amide bond cleavage by BSHs. (**c**) A co-crystal structure of the BSH from the Gram positive gut bacterium *Clostridium perfringens* (strain 13 / type A) and deconjugated tauro-deoxycholic acid (TDCA) (PDB 2BJF) guided our inhibitor design. While hydrophobic interactions orient the bile acid core in the active site, the D-ring side chain and amino acid are exposed to solvent (magenta residues are within 4Å of bile acid, Cys2 is yellow). (**d**) Library of synthesized inhibitors. Electrophilic warheads were appended to the chenodeoxycholic acid bile core in order to create broad-spectrum BSH inhibitors. (**e**) BSH inhibitors previously identified from a high-throughput screen, riboflavin and caffeic acid phenethyl ester (CAPE).

Bile acids, which share a carbon skeleton with steroids, can bind to host receptors, including nuclear hormone receptors (NhR) and G-protein coupled receptors (GPCRs). By acting as either agonists or antagonists for these receptors, bile acids regulate host metabolism, including energy expenditure and glucose and lipid homeostasis,^2,5^ and host immune response, including both innate and adaptive immunity.^6,7^ Dysregulated bile acid metabolism is thought to play causal roles in the pathophysiology of diseases including hypercholesterolemia, obesity, diabetes, cancer, and gallstone formation,^2,8,9^ further highlighting the biological importance of these molecules.

The key reaction in the conversion of primary into secondary bile acids is the hydrolysis of the C24-amide bond of conjugated primary bile acids (Fig. 1a). This enzymatic conversion is performed exclusively by gut bacterial bile salt hydrolase (BSH) enzymes.^1^ BSHs (EC 3.5.1.24) are widespread in human gut bacteria. A recent study identified BSHs in gut species from 117 genera and 12 phyla, including the two dominant gut phyla, Bacteroidetes and Firmicutes, as well as Actinobacteria and Proteobacteria.^10^ A non-toxic, small molecule pan-inhibitor of gut bacterial BSHs would provide a powerful tool to study how bile acids affect host physiology. Such a compound should limit bile acid deconjugation across the vast majority of gut strains without significantly affecting the growth of these bacteria. The use of a pan-inhibitor in vivo would significantly alter bile acid pool composition, shifting the pool toward conjuated bile acids and away from deconjugated bile acids and secondary bile acids (Fig. 1a). This chemical tool would thus allow researchers to investigate previously unanswered questions, including how primary and secondary bile acids differentially affect physiology in a fully colonized host.

Herein, we report the development of a covalent inhibitor of bacterial BSHs using a rational design approach. Importantly, this compound completely inhibits BSH activity in conventional mouse feces, demonstrating its potential utility as a pan-inhibitor of BSHs.

## Results

### Rational design of covalent small molecule inhibitors of bile salt hydrolases

In order to generate potent, long-lasting inhibitors of BSHs, we chose to develop covalent inhibitors of these gut bacterial enzymes. Covalent inhibitors have gained renewed interest in the field of drug discovery due to their ability to inactivate their protein target with a high degree of potency and selectivity even in the presence of large concentrations of native substrate.^11^ The substrates for BSHs, conjugated bile acids, are found in high concentrations in the colon (1-10 mM),^12^ suggesting that covalent inhibition could be an effective strategy for targeting these enzymes. In addition, recently developed irreversible inhibitors of bacterial cutC both block production of trimethylamine N-oxide and display minimal off-target effects.^13^ This work demonstrates that covalent inhibitors of bacterial enzymes can be effective in the gut, thus further validating our approach.

While there is significant divergence in BSH protein sequence across gut strains, all BSHs possess a conserved active site that includes a catalytic cysteine (Cys2).^1,10^ This residue performs the nucleophilic attack on the substrate carbonyl, resulting in amide bond cleavage (Fig. 1b). We reasoned that by designing compounds that targeted this highly conserved Cys residue, we could develop pan-BSH inhibitors. A co-crystal structure of the BSH from Gram positive species *Clostridium perfringens* and the substrate taurodeoxycholic acid (TDCA) showed that hydrophobic interactions held the bile acid core in place and oriented the amide bond toward the conserved cysteine, leaving the amino acid solvent-exposed (Fig. 1c).^14^ Furthermore, purified *C. perfringens* BSH tolerates a large degree of variability in the amino acid side chain, including longer chain conjugates.^15^ These results suggested that the bile acid D-ring side chain was a possible site for incorporation of electrophilic groups into our inhibitors.

Based on this rationale, we designed a small library of potential inhibitors containing both a bile acid core motif to selectively target BSHs and a pendant electrophilic warhead to irreversibly bind the inhibitor to the enzyme (Fig. 1d). While previous literature suggested that BSH enzymes catalyze amide bond cleavage of all conjugated bile acids regardless of the steroidal core,^1,8^ we recently determined that some species from the abundant Gram negative gut bacterial phylum Bacteroidetes cleave C12 = H but not C12 = OH primary bile acids (Fig. 1a).^16^ As our goal was to develop BSH inhibitors that target both Gram negative and Gram positive strains, we decided to utilize the steroidal portion of the human primary bile acid chenodeoxycholic acid (CDCA, C12 = H) as our scaffold.

For the electrophilic trapping groups, we chose warheads that have been successfully deployed in the development of selective and potent protease and kinase inhibitors,^17,18^ including isothiocyanate (**1**),^19^ cyanoacrylate (**2**),^20^ α,β-unsaturated systems (**3** and **4**),^21^ acrylamide (**5**),^22^ and nitrile (**6**).^23^ We also included an inhibitor with an α-fluoromethyl ketone warhead (FMK) (**7**) in our library. Covalent inhibitors with this warhead have been shown to display high potency and selectivity.^24-26^ In contrast to the more electrophilic α-iodo-, α-bromo- and α-chloromethyl ketone warheads, the weak leaving group ability of fluorine renders the FMK warhead less reactive and and hence more selective.^24,26,27^ As a result, FMK-based inhibitors have been shown to result in minimal off-target effects.^24,28^

### Biochemical characterization of BSHs

Following inhibitor synthesis (**Supplementary Note**), we next sought to evaluate the activity of inhibitors **1**-**9** biochemically against both Gram negative and Gram positive BSHs. In particular, we decided to use a selective *Bacteroides* BSH for inhibitor optimization, reasoning that the more limited substrate scope of this enzyme could make it more difficult to target. We heterologously expressed and purified the selective BSH (BT2086) that we had previously identified in *Bacteroides thetaiotaomicron* VPI-5482 (*B. theta*).^16^ We then established kinetic parameters for its hydrolysis of conjugated primary and secondary bile acids using a ninhydrin-based assay.^29^ Consistent with our previous results from *B. theta* cultures, purified *B. theta* BSH displayed a preference for TDCA deconjugation and did not deconjugate TCA (Table 1 and Supplementary Fig. 1).^16^ These results suggest that the enzymatic selectivity observed in *B. theta* culture was due to inherent biochemical properties of the BSH, not to differences in transport or the accessibility of the substrates to the enzyme.

**Table 1:** Kinetic parameters for BSHs from *Bactereroides thetaiotaomicron* (*B. theta*) and *Bifidobacterium longum (B. longum)*.

| BSH source<sup>a</sup> | Substrate<sup>b</sup> | k<sub>cat</sub> (min<sup>-1</sup>) | K<sub>m</sub> (mM) | k<sub>cat</sub>/K<sub>m</sub><br>(min<sup>-1</sup>mM<sup>-1</sup>) |
| --- | --- | --- | --- | --- |
| *B. theta* | TCA<sup>c</sup> | -- | -- | -- |
|  | TUDCA | 15.3 ± 0.8 | 8.2±1.0 | 1.9 ± 0.3 |
|  | TDCA | 12.9 ± 0.6 | 3.4±0.6 | 3.8 ± 0.7 |
|  | TCDCa | 4.3 ± 0.6 | 2.9±1.8 | 4.3 ± 0.9 |
| *B. longum* | TCA | 6.9 ± 0.9 | 8.3±2.5 | 0.8 ± 0.3 |
|  | TUDCA | 0.9 ± 0.2 | 4.1±2.5 | 0.2 ± 0.1 |
|  | TDCA | 3.5 ± 0.1 | 2.3±0.3 | 1.5 ± 0.2 |
|  | TCDCa | 4.6 ± 0.6 | 7.0±2.3 | 0.6 ± 0.2 |

<sup>a</sup>Characterization was performed using ninhydrin reagent and experiments were performed in PBS buffer at pH 7.5 and 37 °C. <sup>b</sup>Conjugated primary and secondary bile acid used as substrates were taurocholic acid (TCA), tauroursodeoxycholic acid (TUDCA), taurodeoxycholic acid (TDCA), taurochenodeoxycholic acid (TCDCa). <sup>c</sup>*B. theta* did not deconjugate TCA.

We also cloned and expressed the known BSH from the Gram positive strain *Bifidobacterium longum* SBT2928 BSH^30^ and determined the kinetic parameters of this enzyme (Table 1 and **Supplementary Fig. 1**). Notably, the *K*_m_ values for all of the recognized substrates are in the low millimolar range, which is approximately the concentration of these bile acids in the gut. While the k_cat_ values for both enzymes are lower than the k_cat_ reported for the BSH from *Lactobacillus salivarius*, the *K*_m_ values for these enzymes are similar to those of previously characterized BSH.^30–32^

### Biochemical evalutation identifies α-FMK compound 7 as lead inhibitor

We next evaluated the ability of the compounds in our library to inhibit *B. theta* and *B. longum* BSH. We also tested riboflavin (**10**) and caffeic acid phenethyl ester (CAPE, **11**), compounds that had been previously identified in a high throughput screen for inhibition of a BSH from a *Lactobacillus salivarius* chicken gut isolate.^33^ (Fig. 1e). To determine the BSH inhibitory activity of these compounds, we pre-incubated the *B. theta* BSH with each inhibitor (100 μM) for 30 minutes and then added a mixture of conjugated bile acids (100 μM final concentration). Hydrolysis of bile acids was monitored by Ultra Performance Liquid Chromatography-Mass Spectrometry (UPLC-MS) over 21 hours. Among the synthesized inhibitors, isothiocyanate (**1**) displayed modest inhibition. Other compounds containing Michael acceptor warheads (inhibitors **2**-**6**) did not inhibit deconjugation. In contrast, incubation with the α-FMK-based inhibitor **7** resulted in almost complete inhibition of the *B. theta* BSH activity for 21 hours (>98%, Fig. 2a and Supplementary Fig. 2). In order to validate that the inhibitory activity of compound **7** was due to the presence of fluorine as a leaving group, we synthesized a methyl ketone analog lacking the fluorine atom (**8**).^28^ This analog did not display BSH inhibition against either recombinant protein or growing *B. theta* cultures, indicating that the α-fluorine group was necessary for activity (Fig. 2a and Supplementary Fig. 3). Riboflavin did not display any inhibitory activity, while CAPE provided only moderate inhibition of *B. theta* BSH.

**Figure 2.**
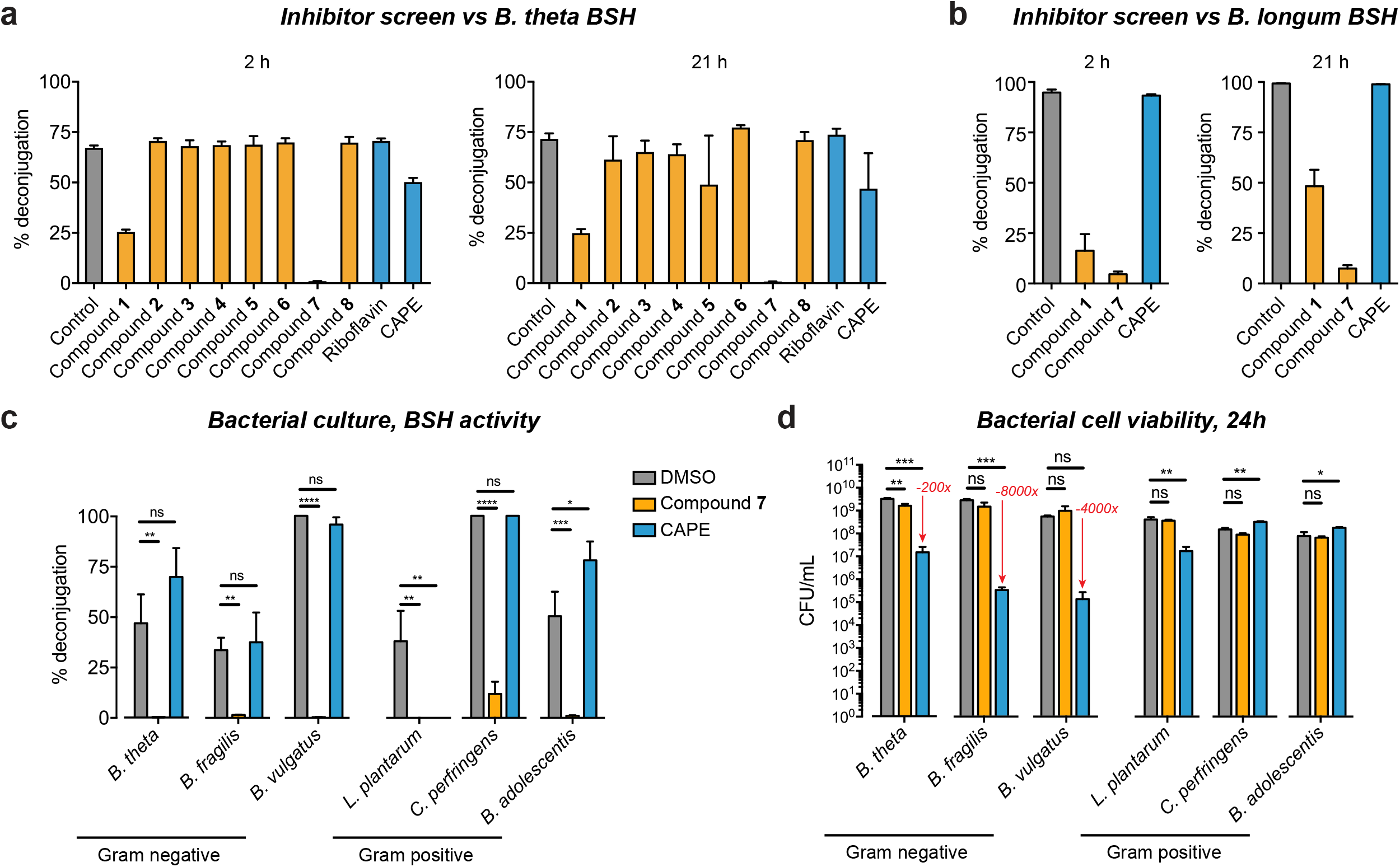
Identification of compound 7 as a potent, non-toxic, broad-spectrum BSH inhibitor. (**a,b**) Screen of inhibitors versus *B. theta* BSH (**a**) and *B. longum* BSH (**b**) showing % deconjugation at 2 and 21 hours. Inhibitor (100 μM) was incubated with 200 nM rBSH for 30 mins followed by addition of taurine-conjugated bile acid substrates (tauro-β-muricholic acid, TβMCA; tauro-cholic acid, TCA; tauro-ursodeoxycholic acid, TUDCA; and tauro-deoxycholic acid, TDCA, 25 μM each). Deconjugation of substrate was followed by UPLC-MS. Assays were performed in biological triplicate. (**c**) Compound **7** inhibited BSH activity in growing cultures of Gram negative (*B. theta* VPI 5482, *Bacteroides fragilis* ATCC 25285, and *Bacteroides vulgatus* ATCC 8482) and Gram positive (*Lactobacillus plantarum* WCFS1, *Clostridium perfringens* ATCC 13124, and *Bifidobacterium adolescentis* L2-32) bacteria. Inhibitor (100 μM of compound **7** or CAPE) and taurine-conjugated bile acid substrates (TβMCA, TCA, TUDCA and TDCA, 25 μM each) were added to bacterial cultures at OD_600_ 0.1. Bacterial cultures were allowed to grow into stationary phase and percent deconjugation at 24h was determined by UPLC-MS. (**d**) Compound **7** is not bactericidal. Bacterial strains were incubated with conjugated bile acids (as described in panel **a**) and compound (100 μM) and plated at 24h to assess the viability of each strain. CAPE decreased the cell viability of the Gram negative strains tested. Red downward arrows indicate fold decrease compared to DMSO control. For (**c**) and (**d**), one-way ANOVA followed by Dunnett’s multiple comparisons test. *p<0.05, **p<0.01, ***p<0.001, ****p<0.0001, ns = not significant. All assays were performed in biological triplicate, and data are presented as mean ± SEM.

We next evaluated the activity of our two most potent inhibitors against *B. theta* BSH, compounds **1** and **7**, as well as CAPE, against the BSH from the Gram positive species *B. longum*. These compounds displayed similar degrees of inhibition of *B. longum* BSH as we had observed against *B. theta* BSH. Compound **7** was again the most active inhibitor, while CAPE was ineffective at inhibiting deconjugation by *B. longum* BSH at all timepoints (Fig. 2b and **Supplementary Fig. 2**). These data indicate that compound **7** is a potent inhibitor of purified BSH protein from both a Gram negative and a Gram positive bacterial strain.

### Compound 7 inhibits BSH activity in growing cultures of gut bacteria

Given that compound **7** displayed activity against purified BSHs, we next sought to evaluate the potency of this inhibitor in growing bacterial cultures. In order to test the scope of BSH inhibition, we included three Gram negative and three Gram positive strains of human gut bacteria known to possess BSH activity (Gram negative, *B. theta*, *Bacteroides fragilis* ATCC 25285, and *Bacteroides vulgatus* ATCC 8482; Gram positive, *Lactobacillus plantarum* WCFS1, *Clostridium perfringens* ATCC 13124, and *Bifidobacterium adolescentis* L2-32) in our screen.^1,16^

Bacterial cultures were diluted to pre-log phase and both inhibitor (100 μM) and a mixture of conjugated bile acids (100 μM final concentration) were added simultaneously. Deconjugation was monitored over 24 hours using UPLC-MS. Strikingly, while all six bacterial strains deconjugated bile acids in the presence of vehicle control, we observed almost no deconjugation in any of the cultures grown in the presence of compound **7** (Fig. 2c). Compound **7** did not significantly affect the cell viability of the majority of the tested strains (Fig. 2d), indicating that the BSH inhibition observed was not due to bactericidal activity. To quantify the potency of compound **7**, we determined that the IC_50_ values of this inhibitor against the Gram negative strain *B. theta* and the Gram positive strain *B. adolescentis* were 912 nM and 240 nM, respectively (**Supplementary Fig. 4**). Taken together, these results indicate that compound **7** is a potent, broad-spectrum inhibitor of BSHs.

In contrast, we observed no inhibition of deconjugation over the course of 21 hours in five out of the six bacterial strains grown in the presence of CAPE (100 μM) (Fig. 2c). CAPE was found to inhibit deconjugation in *L. plantarum*, suggesting that this compound may inhibit BSHs from *Lactobacilli* but is not a broad-spectrum BSH inhibitor. Moreover, CAPE inhibited the cell viability of all three Gram negative bacterial strains tested (Fig. 2d, ~200-fold, ~8000-fold, and ~4000-fold decreases in 24h CFU/mL compared to DMSO control for *B. theta*, *B. fragilis*, and *B. vulgatus*, respectively). These results suggest that the dominant effect of CAPE on Gram negative bacteria is not inhibition of BSH activity but rather inhibition of growth.

Finally, to evaluate our hypothesis that C12 = OH compounds would not be effective broad-spectrum inhibitors, we synthesized a potential inhibitor in which we appended the α-FMK warhead to a C12 = OH bile acid core, cholic acid (compound **9**, Fig. 1d). Compound **9** displayed significantly reduced ability to inhibit BSH deconjugation in growing *B. theta* cultures compared to compound **7** (**Supplementary Fig. 3**). These results support the hypothesis that bile acid core structure, specifically C12 substitution, affects the ability of our probes to inhibit selective BSH. In addition, these results suggest that the α-FMK warhead is not broadly reactive but rather requires suitable positioning within the active site, a hypothesis that we further investigated using mass spectrometry and crystallography studies.

### Compound 7 covalently modifies the catalytic cysteine residue of BSH

With the potency of compound **7** established, we next investigated its mechanism of inhibition. The *B. theta* BSH contains two cysteine residues, Cys2 and Cys67. Analysis of an apo crystal structure of this enzyme revealed that both the cysteine residues are pointed towards the active site, indicating either residue could be a potential site for covalent modification (PDB 3HBC). To confirm that compound **7** is a covalent inhibitor that modifies Cys2, we incubated purified BSH enzyme with an excess of this molecule. Analysis by mass spectrometry revealed a mass shift consistent with the addition of a single molecule of compound **7**, confirming formation of a covalent bond (Fig. 3a). Subsequent top-down mass spectrometry analysis identified Cys2 as the modified residue (Fig. 3b).

**Figure 3.**
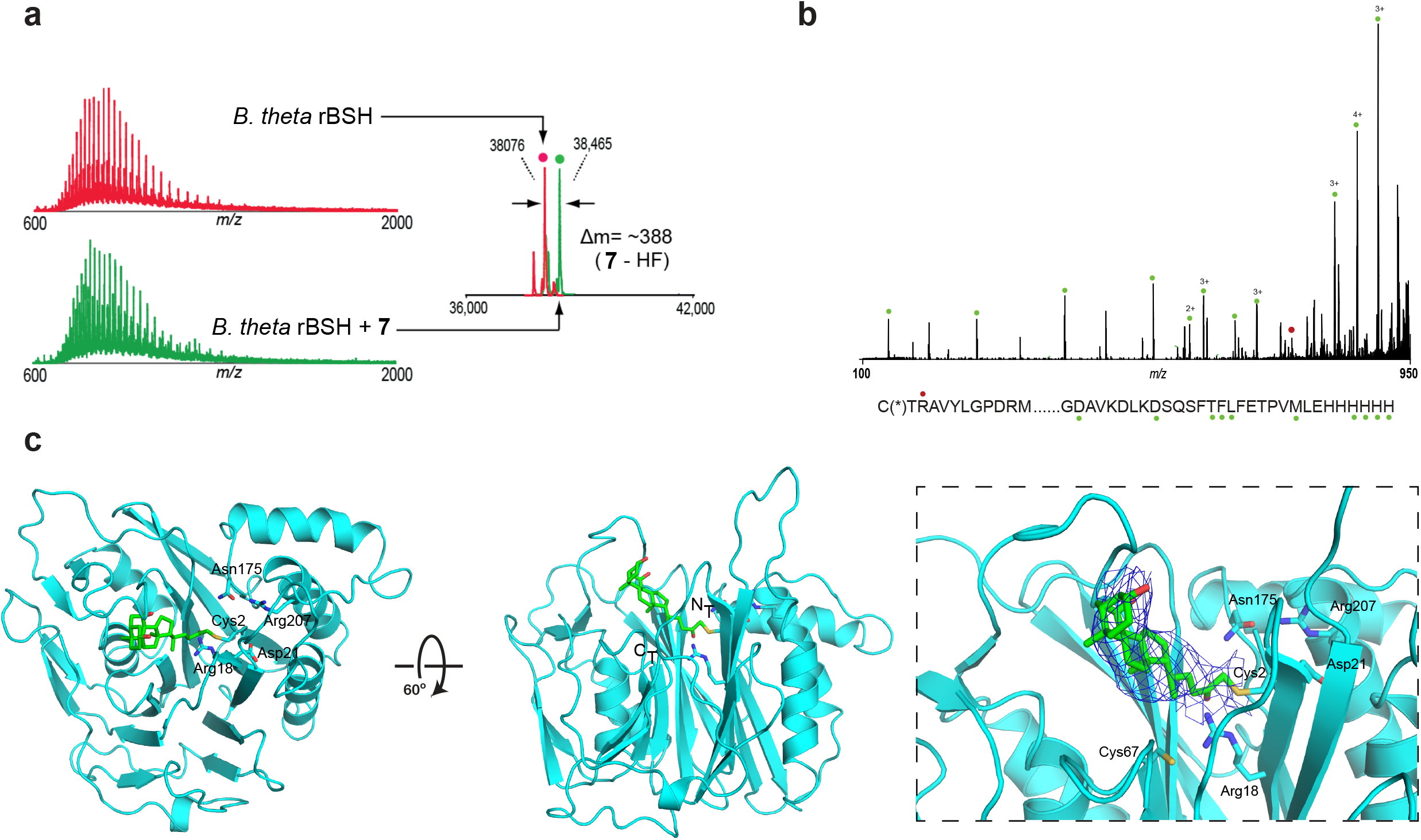
Compound 7 covalently modifies *B. theta* BSH at the active site cysteine residue. (**a,b**) Mass spectrometry revealed that compound **7** monolabels *B. theta* BSH. (**a**) Mass spectra (left) and zero-charge mass spectra (right) of BSH treated with DMSO (top, trace in red) or 10-fold excess of inhibitor compound **7** for 2 h (bottom, trace in green). A shift in mass of 388 Da is consistent with covalent modification of BSH with a loss of HF. (**b**) Top-down MS/MS of BSH treated with 10 fold excess of compound **7**. Ions of type c and z are indicated with red and green glyphs respectively. Ion c3 indicates that modification is on the N-terminus Cys2 residue. (**c**) X-ray structure of compound **7** bound to *B. theta* BSH. The BSH (cyan) is shown in ribbon representation, with indicated side chains (cyan, with heteroatoms in CPK colors) rendered as sticks. Compound **7** (green, with heteroatoms in CPK colors) is rendered in stick form. There is electron density visible at the active site of one of the four subunits in the asymmetric unit, consistent with the conclusion that the inhibitor is covalently attached to Cys2. Panel C was prepared using PYMOL software (Schroedinger).

In order to elucidate the position of the bound inhibitor and guide further inhibitor design, we determined the structure of the *B. theta* BSH, first in its apo form to 2.7 Å resolution and then covalently bound to compound **7** to 3.5 Å resolution (**Supplementary Table 1**) (PDB XXX). The structure of the BSH-inhibitor complex contains four copies of the protein in the asymmetric unit. The electron density map is best resolved in two of the four subunits, and electron density is clearly visible for the inhibitor in one of these subunits covalently attached to Cys2 (Fig. 3c). Comparison with the apo structure also suggests that there is a loop (residues 127-138) which repositions to clasp the inhibitor in the active site in a solvent-exposed channel (**Supplementary Fig. 5**). Taken together, these data indicate that compound **7** selectively labels the *B. theta* BSH at the catalytic cysteine residue. Furthermore, the co-crystal structure reveals that the C3-hydroxyl group is solvent-accessible, suggesting that this site might be amenable to further modification (**Supplementary Fig. 5**).

### Compound 7 displays minimal off-target effects

While covalent inhibitors have been shown to be highly potent, concerns have been raised that non-specific reactivity of these compounds could result in acute toxicity.^11^ Our inhibitors were designed to contain a bile acid core in order to increase selectivity of these compounds for BSHs. However, bile acids are known to be ligands for host nuclear hormone receptors (NhR) and G protein-coupled receptors (GPCR), in particular, the farnesoid X receptor (FXR) and the G protein-coupled bile acid receptor 1 (GPBAR1, or TGR5).^2^ In order to determine whether compound **7** could act as a ligand for FXR, we performed an in vitro coactivator recruitment assay.^34^ While the known FXR agonist GW4064 showed a clear dose-dependent increase in the binding of the co-activator peptide SRC2-2 to FXR (EC_50_=50 nM), the binding of SRC2-2 to FXR did not increase in the presence of compound **7**, suggesting that this inhibitor does not activate FXR (Fig. 4a). In the presence of GW4064, compound **7** did not display a dose-dependent curve, indicating that compound **7** does not possess FXR antagonist activity at physiologically relevant concentrations (**Supplementary Fig. 6**). Next, we evaluated the effect of compound **7** on TGR5 activation in a human intestinal cell line (Caco-2). Compound **7** did not agonize TGR5 over the range of concentrations tested (Fig. 4b). In addition, compound **7** did not antagonize TGR5 in the presence of known TGR5 agonist LCA (10 μM) (**Supplementary Fig. 6**). These results suggest that compound **7** does not induce off-target effects on either of these critical host receptors.

**Figure 4.**
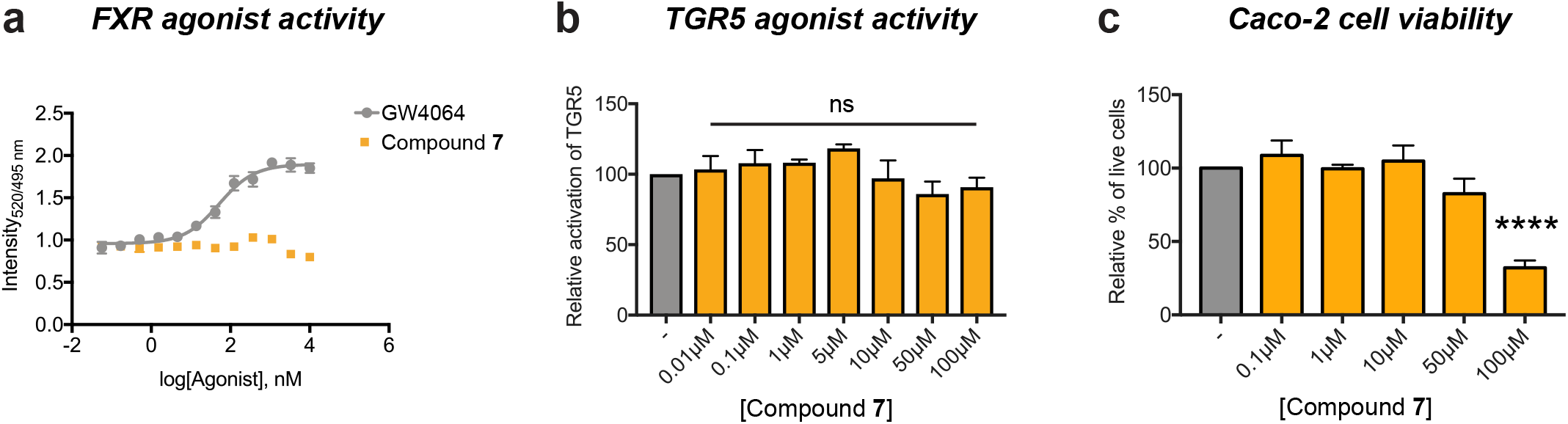
Compound 7 exhibits minimal off-target effects. (**a**) Compound **7** is not an farnesoid X receptor (FXR) agonist as determined by an FXR coactivator recruitment assay. n=4 biological replicates per concentration. (**b**) Compound **7** is not a G protein-coupled bile acid receptor (GPBAR1 / TGR5) agonist. Endogenous TGR5 agonist activity was measured by incubating Caco-2 cells with varying concentrations of compound **7** overnight. (**c**) Compound **7** did not display toxicity toward Caco-2 cells up to a concentration of 50 μM. For (**b**) and (**c**), n≥3 biological replicates per concentration, one-way ANOVA followed by Dunnett’s multiple comparisons test, ns = not significant, ****p<0.0001. All data are presented as mean ± SEM.

In addition to their effects on host receptors, bile acids are known to be toxic to cells due to their detergent properties.^1,35^ Because the expected in vivo area of action of compound **7** is the lower gut, we tested the toxicity of this compound against human intestinal cells (Caco-2). No resultant toxicity was observed when these cells were incubated with up to 50 μM of compound **7** (Fig. 4c). Because the IC_50_ values of compound **7** against bacterial BSHs range from 240 to 912 nM, these results suggest that it should be possible to achieve an effective in vivo dose at a concentration that will not result in toxicity to intestinal cells. Taken together, our results indicate that compound **7** is both non-toxic and selective for bacterial BSHs over potential host targets.

### Compound 7 inhibits BSH activity in conventional mouse feces

While we were able to demonstrate the potency of compound **7** against growing cultures of six different strains of gut bacteria, there are hundreds of bacterial species in the human gut.^36^ Previous literature had reported significant BSH activity in mouse feces.^37^ To further assess whether compound **7** is a pan-inhibitor of BSH, we tested its activity in resuspended feces from conventional mice. Compounds **1**, **7**, and CAPE (20 μM) were added to a fecal suspension in buffer. After 30 minutes, the deuterated substrate GCDCA-d4 was added, and deconjugation was quantified after 18 hours using UPLC-MS (Fig. 5a). Strikingly, we observed that incubation with compound **7** completely inhibited the BSH activity in feces (Fig. 5b). Consistent with our in vitro results, CAPE provided no inhibition of BSH activity in mouse feces. These results demonstrate that compound **7** is a potent, pan-inhibitor of gut bacterial BSH activity.

**Figure 5.**
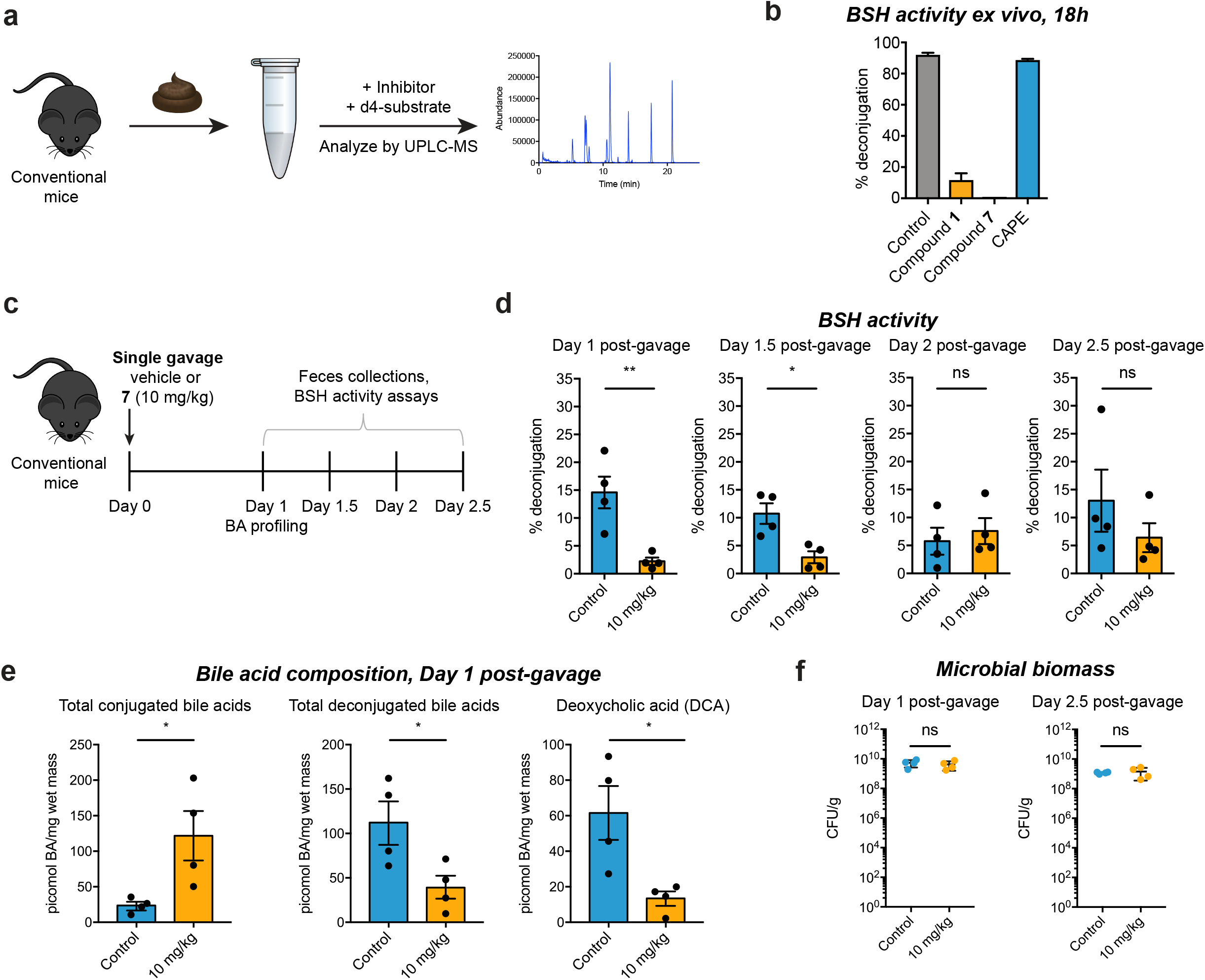
Compound 7 inhibits BSH activity ex vivo and in vivo. (**a**) Fecal BSH activity assay design. Freshly collected feces from conventional mice were resuspended in PBS (1 mg/mL) and incubated with 20 μM of inhibitor (Compound **1**, **7** or CAPE) for 30 mins. Glycochenodeoxycholic acid-*d*4 (GCDCA-*d*4, 100 μM) was added as a substrate and deconjugation was determined by UPLC-MS after 18 hours. (**b**) Compound **7** inhibited BSH activity in a fecal slurry. Assays were performed in biological triplicate. (**c**-**e**) Treatment of conventional mice with a single dose of compound **7** resulted in recoverable inhibition of BSH activity and a shift toward conjugated bile acids. n=4 mice per group, Welch’s t test, *p<0.05, **p<0.01, ns = not significant. (**c**) Design of in vivo BSH inhibition experiment. Adult male C57BL/6 mice were gavaged with a single dose of compound **7** (10 mg/kg) or vehicle control. (**d**) BSH activity was measured in half-day increments starting 1 day post-gavage. Resuspended fresh feces from inhibitor-or vehicle-treated groups were incubated with substrate (GCDCA-*d*4, 100 μM) for 25 min and deconjugation was quantified by UPLC-MS. (**e**) Fecal bile acid composition 1 day post-gavage. Deconjugated bile acids, including the secondary bile acid deoxycholic acid (DCA), were decreased in the inhibitor-treated group. (**f**) Microbial biomass did not differ between the inhibitor- and vehicle-treated groups 1 day or 2.5 days post-gavage. n=4 mice per group, Mann-Whitney test. All data are presented as mean ± SEM.

### Single dose of compound 7 inhibits BSH activity in conventional mice

Having established the potency of compound **7** in vitro, we next sought to evaluate the activity of this inhibitor in conventional mice. C57BL/6 mice were gavaged with a single dose of either compound **7** (10 mg/kg) or vehicle control, and BSH activity was monitored over time in half-day increments until 2.5 days post-gavage (Fig. 5c). We predicted that if compound **7** was active in vivo, we would observe an initial decrease in BSH activity followed by recovery of activity. Gratifyingly, we observed this expected effect. One day and 1.5 days post-gavage, we noted a significant decrease in BSH activity in feces, while at subsequent timepoints, BSH activity recovered (Fig. 5d). Consistent with these results, we observed a significant increase in conjugated bile acids and a decrease in deconjugated bile acids 1 day-post gavage. Notably, we observed a decrease in the deconjugated secondary bile acid deoxycholic acid (DCA) at this timepoint (Fig. 5e). In agreement with our in vitro bacterial cell viability results, we did not observe a significant decrease in bacterial biomass following gavage with compound **7** (Fig. 5f and **Supplementary Fig. 7**). Taken together, our results indicate that compound **7** can inhibit gut bacterial BSH activity and modulate the bile acid pool in vivo while not significantly inhibiting overall growth of the gut bacterial community.

## Discussion

In order to uncover the effects that bacterial metabolites have on host health, tools are needed that selectively control the levels of these compounds in fully colonized animals. In this work, we report the development of such a chemical tool, a potent, selective, pan-inhibitor of gut bacterial BSHs. We identified a lead inhibitor, compound **7**, that effectively inhibits deconjugation by purified BSH protein, growing cultures of both BSH-containing Gram negative and Gram positive human gut strains, and resuspended conventional mouse feces. We then showed that a single dose of compound **7** administered to conventional mice reduces BSH activity and predictably shifts the in vivo bile acid pool. Importantly, we found that compound **7** does not significantly affect the viability of these bacteria.

Our results suggest that compound **7** or derivatives thereof can be used as tools to study the biological roles of primary and secondary bile acids in conventional animals, including both wild-type and knock-out mouse strains. These investigations would broaden our understanding of how bile acids affect host immune and metabolic systems and reveal how these metabolites affect the composition and biogeography of the gut bacterial community.

In particular, BSH inhibitors could be used to better understand the effects of bile acids on host metabolism. Previous studies have reported conflicting results about how altering BSH activity in vivo may affect host metabolic responses.^37,39^ In these studies, the introduction into the gut of an exogenous bacterial strain overexpressing a heterologous gene or the use of a molecule that may exert metabolic effects through a BSH-independent mechanism do not permit evaluation of how BSH activity affects metabolism in the native host-bacterial system. Moreover, in previous work, we showed that deleting the BSH-encoding gene from *B. theta* resulted in decreased weight gain and a decreased respiratory exchange ratio in mice colonized with this bacterium compared to mice colonized with the *B. theta* wild-type strain.^16^ However, these experiments were performed in monocolonized germ-free mice and do not reveal how limiting activity of all BSHs will affect the metabolism of conventional animals. Use of a non-toxic, pan-BSH inhibitor would enable investigation of how BSH activity directly affects metabolism in fully colonized hosts.

In addition, BSH inhibitors could enable investigation of how bile acids affect host immune response in the context of liver cancer. A recent study proposed a causal connection between the conversion of primary to secondary bile acids and a decrease in a tumor-suppressive environment in the liver mediated by the accumulation of beneficial NKT cells.^40^ Use of a BSH inhibitor in mouse models of liver cancer could further test this hypothesis by shifting the endogenous in vivo bile acid pool toward primary bile acids without significantly perturbing the enterohepatic system and the microbial community.

Finally, because BSH inhibitors would remove secondary bile acids from the bile acid pool, such inhibitors could be used in chemical complementation experiments in which individual secondary bile acids were reintroduced through feeding. These studies would reveal how specific secondary bile acids affect host physiology and bacterial community composition in conventional animals. Looking ahead, if use of BSH inhibitors in vivo beneficially affects host physiology, by limiting liver tumor growth, for example, or decreasing liver steatosis, these compounds could be developed as novel drug candidates. In this way, development of mechanism-based, non-toxic chemical probes for bacterial enzymes could lay the groundwork for potential therapeutic agents that target the microbiome.^13,38^

## Acknowledgements

This research was supported National Institutes of Health (NIH) grant R35 GM128618 (A.S.D), R35 CA220340 (S.C.B.), R01 CA222218 (J.A.M.), an Innovation Award from the Center for Microbiome Informatics and Therapeutics at MIT (A.S.D), a grant from Harvard Digestive Diseases Center (supported by NIH grant 5P30DK034854-32) (A.S.D), a Karin Grunebaum Cancer Research Foundation Faculty Research Fellowship (A.S.D), a John and Virginia Kaneb Fellowship (A.S.D), a Quadrangle Fund for the Advancement and Seeding of Translational Research at Harvard Medical School (Q-FASTR) grant (A.S.D), and an HMS Dean’s Innovation Grant in the Basic and Social Sciences (A.S.D). L.Y. and S.N.C. acknowledge a Wellington Postdoctoral Fellowship and an American Heart Association Postdoctoral Fellowship, respectively. M.D.M. acknowledges an NSF Graduate Research Fellowship (DGE1745303). D.R. is supported by a grant from the Swiss National Science Foundation. We are indebted to Nathanael Gray, David Scott, John M. Hatcher, Jinhua Wang, Jon Clardy, Matthew Henke, and members of the Clardy group for helpful discussions. We thank the ICCB-Longwood Screening Facility for use of their fluorescent plate reader.

## Author Contributions

A.A.A and A.S.D conceived the project and designed the experiments. A.A.A. performed most of the experiments. T.C.S. and S.C.B. performed the crystallization studies. S.B.F. and J.A.M. performed the mass spectrometry studies. D. R. and A.S.B. performed the in vivo experiment and provided fresh mouse feces. M.D.M. purified and performed experiments with *B. longum* BSH. L.Y. performed the in vitro FXR assays and provided help with experiments. S.N.C. performed the cell culture assays. S.N. assisted with bacterial culture experiments. A.A.A. and A.S.D wrote the manuscript. All authors edited and contributed to the critical review of the manuscript.

### Online Methods

#### Reagents

All bile acids were commercially purchased from Steraloids Inc. and Sigma Aldrich. Stock solutions of all bile acids and inhibitors were prepared in molecular biology grade DMSO (Sigma Aldrich) at 1000X concentrations. These bile acid stock solutions were also used for establishing standard curves. Solvents used for preparing UPLC-MS samples were HPLC grade. New biological materials reported here are available from the authors upon request.

#### Chemical Synthesis

See supplementary information for detailed procedures and characterization data of all compounds.

#### Bacterial Culturing

All bacterial strains were cultured at 37 °C in Cullen-Haiser Gut (CHG) media supplemented with hemin and Vitamin K. All strains were grown under anaerobic conditions in a anaerobic chamber (Coy Lab Products Airlock) with a gas mix of 5% hydrogen and 20% carbon dioxide nitrogen. *Escherichia coli* was grown aerobically at 37 °C in LB medium supplemented with ampicillin to select for the pET21b plasmid.

#### Protein Expression and Purification

##### *B. thetaiotaomicron* rBSH

The gene encoding BT2086 (without the leader sequence) was codon-optimized for *E. coli* and cloned into pET-21b(+) vector containing a C-terminal His_6_ tag. The expression plasmid was then transformed into BL21(DE3)pLysS *Escherichia coli* (New England Biolabs) cells under ampicillin selection. Overnight starter cultures were grown in LB media with ampicillin (50 μg/mL) and diluted 1:1000 in fresh LB media with ampicillin and grown at 37 °C. Expression was induced at an OD_600_ of 0.6-0.7 by the addition of 1 mM isopropyl-1-thio-D-galactopyranoside (IPTG) and further incubated at 18 °C overnight. The cells were pelleted by centrifugation at 7,000g for 20 mins at 4°C. The pelleted cells were then resuspended in buffer (50 mM sodium phosphate at pH 7.5, 300 mM NaCl, 5% glycerol) along with 20 mM imidazole, 1mM phenylmethylsulfonyl fluoride (PMSF), and 0.25 mM tris(2-carboxyethyl)phosphine hydrochloride (TCEP). The resuspended cells were sonicated and then pelleted by centrifugation at 16,000g for 20 mins at 4 °C. The supernatant was then mixed with pre-formed Ni-NTA for 45 mins at 4 °C. The slurry was loaded onto a column and allowed to drain under gravity. The nickel-bound protein was eluted with buffer (50 mM sodium phosphate at pH 7.5, 300 mM NaCl, 0.25 mM TCEP, 5% glycerol) containing gradually increasing concentration of imidazole. Collected fractions were tested for purity by SDS-PAGE. The pure fractions were combined and concentrated followed by dialysis using the storage buffer (50 mM sodium phosphate at pH 7.5, 300 mM NaCl, 0.25 mM TCEP and 5% glycerol).

For crystallization purposes, the protein was further purified using S200 size exclusion column (from GE) on a BioRad FPLC in 50 mM tris(hydroxymethyl)aminomethane buffer with 300 mM NaCl, 0.25 mM TCEP and 5% glycerol at pH 7.5.

##### *B. adolescentis* rBSH

Recombinant BSH from *B. adolescentis* SBT2928 was expressed and purified as above, except 0.25 mM IPTG was used for protein expression and 1 mM TCEP for protein purification.

##### Enzyme Kinetics

The enzyme was characterized using a modified BSH activity assay.^29^ To 144.8 μL reaction buffer (PBS at pH 7.5 containing 10 mM TCEP and 5% glycerol), 35.2 μL of recombinant BSH was added as a solution in PBS at pH 7.5 containing 0.25 mM TCEP and 5% glycerol to afford a final concentration of 6.2 μM and 7.0 μM for *B. theta* BSH and *B. longum* BSH, respectively. This solution was preheated to 37 °C in a water bath. 20 μL of a conjugated bile acid in DMSO at appropriate concentration was preheated to 37 °C in a water bath and added to the above solution. At every time interval 15 μL of the mixture was quenched with 15 μL of 15% trichloroacetic acid. The cloudy solution was then centrifuged at 4,200 x g for 15 mins. 10 μL of the supernatant was added to 190 μL of ninhydrin mix (15 mL of 1% [wt/vol] ninhydrin in 0.5 M sodium citrate at pH 5.5, 36mL glycerol and 6 mL 0.5 M sodium citrate buffer at pH 5.5) and the mixture was then heated to 100 °C in a BioRad thermocycler for 18 mins. The obtained solution was cooled at 4 °C for 20 mins and absorbance was measured at 570 nm using a spectrophotometer (Molecular Devices).

##### Inhibitor Screen Using rBSHs

200 nM of BSH was incubated with 100 μM of inhibitor at 37 °C for 30 mins in 3 mL PBS buffer containing 0.25 mM TCEP and 5% glycerol at pH 7.5. 100 μM Bile acid pool consisting of TCA, TβMCA, TUDCA and TDCA (25 μM each) was added to the above solution and incubated at 37 °C. At timepoint intervals, 1 mL of the above buffer solution was acidified to pH = 1 using 6M HCl and extracted twice with 1mL ethyl acetate. The combined organic layers were then dried using a Biotage TurboVap LV. The dried extracts were resuspended in 1:1 methanol:water. Samples were analyzed by UPLC-MS (Agilent Technologies 1290 Infinity II UPLC system coupled online to an Agilent Technologies 765 6120 Quadrupole LC/MS spectrometer in negative electrospray mode) using a previously published bile acid analysis method.^16^

##### Inhibitor Screen in Bacteria

Overnight starter cultures of bacteria were diluted to OD_600_ of 0.1 in 4 mL CHG media containing 100 μM of the taurine conjugated bile acid pool consisting of TCA, TβMCA, TDCA and TUDCA (25 μM each) and various inhibitors at a concentration of 100 μM. These cultures were then allowed to grow anaerobically at 37 °C. After 24 hours, serial dilutions on BHI (Brain Heart Infusion) agar supplemented with vitamin K and hemin were performed to determine cell viability (CFU/mL), and the cultures were extracted and analyzed as per the method described in “Inhibitor Screen Using rBSHs”.

##### Determination of IC_50_ Values of Compound 7

Overnight starter cultures of *B. theta* and *B. adolescentis* were diluted to an OD_600_ of 0.1 in 2 mL fresh CHG containing 100 μM TUDCA or TDCA, respectively, and inhibitor **7** at various concentrations. *B. theta* and *B. adolescentis* deconjugated TUDCA and TDCA, respectively, to the greatest extent of any of the conjugated substrates in the Inhibitor Screen in Bacteria assay, and therefore these substrates were used to determine IC_50_ values. Cultures were then allowed to grow anaerobically at 37 °C for 24h (*B. adolescentis*) or 48h (*B. theta*). Longer incubation time was required for *B. theta* because for this bacterium, significant BSH activity was only observed during stationary phase. Cultures were extracted and analyzed as per the method described in “Inhibitor screen using rBSH”.

##### Crystallization, Data Collection, and Structure Determination

Crystals of BSH and BSH in complex with compound **7** were grown in 24-well format hanging drops at room temperature. BSH crystals (5 mg/mL) grew from micro seeding after 3 days in 42% tacimate 100 mM Tris pH 7.4. The BSH-compound **7** complex (5.0 mg/mL) crystals grew after 5 d in 21% PEG 3350 and 100 mM X Sodium citrate tribasic dihydrate pH 5.0. Crystals were cryoprotected by supplementing the mother liquor with 10% 2-methyl-2,4-pentanediol (v/v). Individual crystals were flash frozen in liquid nitrogen and stored until data collection. Data collection was performed at Advanced Photon Source NE-CAT beamline 24 ID-C. Diffraction images were processed and scaled using XDS.^41^ To obtain phases for the apo BSH structure, molecular replacement was performed in Phenix with Phaser^42^ choloylgylcine hydrolase from *B. thetaiotaomicron*, PDB 3HBC as the search model. Iterative model building and reciprocal space refinement was performed in COOT^43^ and phenix.refine,^44^ respectively. Reciprocal space refinement used reciprocal space optimization of xyz coordinates, individual atomic B-factors, NCS restraints, optimization for X-ray/stereochemistry weights, and optimization for X-ray/ADP weights. The BSH-compound **7** structure was phased using molecular replacement for all four copies in the asymmetric unit with the apo BSH as a search model. Iterative model building and refinement for the BSH-compound **7** was similar to the apo BSH structure with changes to grouping atomic B-factors and the addition of an applied twinlaw of *k h –l*. Model quality for both structures was evaluated using composite omit density maps. In final cycles of model building, NCS restraints were removed. Final model quality was assessed using MolProbity.^45^ All crystallographic data processing, refinement, and analysis software was compiled and supported by the SBGrid Consortium.^46^ Data acquisition and refinement statistics are presented in **Supplementary Table 1**. Figures were prepared using Pymol (Schrödinger).

##### Mass Spectrometry Analysis

BSH protein was incubated with DMSO or a 10-fold molar excess of inhibitor **7** for 2 hours at room temperature. Reactions were then analyzed by LC-MS using a Shimadzu LC and autosampler system (Shimadzu, Marlborough, MA) interfaced to an LTQ ion trap mass spectrometer (ThermoFisher Scientific, San Jose, CA). Protein (5 ug) was injected onto a self-packed RP column (5 cm POROS 50R2, Applied Biosystems, Foster City, CA), desalted for 4 minutes with 100%A, eluted with a ballistic gradient (0-100% B in 1 minute; A= 0.2 M acetic acid in water, B= 0.2M acetic acid in acetonitrile), and introduced to the mass spectrometer by ESI (spray voltage = 4.5 kV). The mass spectrometer was programmed to collect full scan mass spectra in profile mode (*m/z* 300-2000). Mass spectra were deconvoluted using MagTran version 1.03b2.^47^

To determine the site of modification, compound **7** modified protein was analyzed as described above, except that the LC system was interfaced to an Orbitrap Lumos Mass Spectrometer (ThermoFisher Scientific). The mass spectrometer was programmed to perform continuous cycles consisting of 1 MS scan (*m/z* 300-2000, profile mode, electron multiplier detection) followed by ETD MS/MS scans targeting the +41 charge state precursor of compound **7** modified protein (ETD reagent target = 200 ms, image current detection at 60K resolution, target value=2E6, ETD reaction time= 100 or 200 ms). Ion assignments were performed using mzStudio software.^48^

##### Effect of Compound 7 on FXR

LanthaScreen TR-FRET Coactivator Assay (Invitrogen, Carlsbad, CA) was used to test the effect of compound **7** on FXR. Test compounds were diluted in DMSO, and assays were run per the manufacturer’s instructions. Known FXR agonist GW4064 (Sigma, G5172) was used as a positive control (agonism assay) or added at its EC_50_ (50.3 nM, measured in this assay) (antagonism assay). Following 1 hour incubation at room temperature, the 520/495 TR-FRET ratio was measured with a PerkinElmer Envision fluorescent plate reader using the following filter set: excitation 340 nm, emission 495 nm, and emission 520 nm. A 100 μsec delay followed by a 200 μsec integration time was used to collect the time-resolved signal.

##### Cell Culture

Caco-2 cells were obtained from American Type Culture Collection (Manassas, VA). Caco-2 cells were maintained in Minimum Essential Medium (MEM) supplemented with GlutaMAX and Earle’s Salts (Gibco, Life Technologies, UK). All cell culture media were supplemented with 10% fetal bovine serum (FBS), 100 units/ml penicillin, and 100μg/ml streptomycin (GenClone). Cells were grown in FBS- and antibiotic-supplemented ‘complete’ media at 37°C in an atmosphere of 5% CO_2_.

##### Cell Viability Assay

Caco-2 cells were treated with compound **7** diluted in DMSO in complete MEM media. The concentration of DMSO was kept constant and used as a negative control. Cells were incubated with compound **7** overnight at 37°C in an atmosphere of 5% CO_2_. The next day, cells were treated with 0.25% trypsin in HBSS (GenClone) for 10 min at 37°C. Cell viability was measured in Countess II automated cell counter (Invitrogen). Percentage relative viability was calculated compared to DMSO control.

##### Plasmids and Transient Transfections

For luciferase reporter assays, vectors expressing human reporter constructs were used. The pGL4.29[luc2P/CRE/Hygro] plasmid (Promega Corporation) was transiently transfected in Caco-2 cells at a concentration of 2 μg/ml of media each for studying TGR5 activation respectively. The pGL4.74[hRluc/CMV] plasmid (Promega Corporation) was used as a transfection efficiency control at a concentration of 0.05 μg/ml of media. All plasmids were transfected using Opti-MEM (Gibco) and Lipofectamine 2000 (Invitrogen, Life Technologies, Grand Island, NY, USA) according to manufacturer’s instructions. Plasmid transfections were performed in antibiotic-free MEM media with 10% FBS. After overnight incubation, compound **7** and/or bile acids were added in complete media. Compound **7** and/or bile acids were diluted in DMSO and the concentration of DMSO was kept constant. 10 μM of LCA was added along with compound **7** to study TGR5 antagonism and incubated overnight. Cells were harvested the next day for luciferase assay.

##### Luciferase Reporter Assay

Luminescence was measured using the Dual-Luciferase Reporter Assay System (Promega Corporation) according to manufacturer’s instructions. Cells were washed gently with PBS and lysed in PLB from the kit. Matrigel-attached cells were scraped in PLB. Luminescence was measured using a SpectraMax M5 plate reader (Molecular Devices, San Jose, CA) at the ICCB-Longwood Screening Facility at HMS. Luminescence was normalized to *Renilla* luciferase activity and percentage relative luminescence was calculated compared to DMSO control.

##### Screen of Inhibitors in Conventional Mouse Feces

BSH activity in fecal pellets were quantified using a modified version of a published method.^49^ Fecal pellets (approximately 10-20 mg) were broken into fine paticles in buffer (10% PBS, 90% sodium acetate at pH 5.2) to obtain a concentration of 1 mg/mL. 20 μM of inhibitors and CAPE were added to the fecal slurry and the mixture was incubated at 37 °C for 30 mins. 100 μM glycochenodeoxycholic acid-d4 (GCDCA-d4) was then added to the mixture and the mixture was incubated at 37 °C for 18 hours. The tubes were then frozen in dry ice for 5 mins and upon thawing were diluted with an equal volume of methanol. The slurry was then centrifuged at 12,500g for 10 mins. The supernatant was decanted into a clean Eppendorf tube and centrifuged again. The supernatant was then transferred to MS vials and analyzed by UPLC-MS using a previously published method.^16^

##### Animal Studies

C57BL/6 mice obtained from Jackson laboratories were maintained under a strict 12 h/12h light/dark cycle and a constant temperature (21 ± 1 °C) and humidity (55–65%). All experiments were conducted on 8–9 week old male mice. The mice were maintained on a standard chow diet (LabDiet, no. 5053) for the duration of the experiment. Mice were split into two groups of four mice each. The vehicle group were gavaged with 200 μL of corn oil containing 5% DMSO. The experimental group were gavaged with 200 μL of corn oil containing compound **7** at a concentration of 1.25 mg /mL. For the fecal pellet collection, each mouse was transferred to a temporary cage for a few minutes until it defecated. Once these fresh fecal pellets were collected, the mice were transferred back to their home cages. All experiments involving mice were performed using IACUC approved by the Beth Israel Deaconess Medical Center.

##### BSH Activity in Feces

BSH activity in fecal pellets were quantified using a modified version of a published method.^49^ Fecal pellets (approximately 10-20 mg) were suspended in buffer (10% PBS, 90% sodium acetate at pH 5.2) containing 100 μM glycochenodeoxycholic acid-d4 (GCDCA-d4) to obtain a concentration of 20 mg/mL. The fecal pellets were broken into fine particles and the mixture was incubated at 37 °C for 25 mins. Samples were analyzed as per the method described in “Screen of Inhibitors in Conventional Mouse Feces.

##### Quantification of Fecal Bile Acids

Bile acid extraction and analysis of pre-massed fecal pellets (~10-20 mg/sample) was performed using a previously published method.^16^

##### Determination of Microbial Biomass

Frozen fecal pellets were used to determine colony forming units (CFU/g). Feces were suspended in PBS buffer in an anaerobic chamber. Serial dilutions were plated on CHG agar plates and incubated at 37 °C.

## References

1. Ridlon, J. M., Kang, D.-J. & Hylemon, P. B. Bile salt biotransformations by human intestinal bacteria. J. Lipid Res. 47, 241–259 (2006).

2. Fiorucci, S. & Distrutti, E. Bile Acid-Activated Receptors, Intestinal Microbiota, and the Treatment of Metabolic Disorders. Trends Mol Med 21, 702–714 (2015).

3. Setchell, K. D., Lawson, A. M., Tanida, N. & Sjövall, J. General methods for the analysis of metabolic profiles of bile acids and related compounds in feces. J. Lipid Res. 24, 1085–1100 (1983).

4. Hamilton, J. P. et al. Human cecal bile acids: concentration and spectrum. Am. J. Physiol. Gastrointest. Liver Physiol. 293, G256–G263 (2007).

5. Modica, S., Gadaleta, R. M. & Moschetta, A. Deciphering the nuclear bile acid receptor FXR paradigm. Nucl Recept Signal 8, e005 (2010).

6. Vavassori, P., Mencarelli, A., Renga, B., Distrutti, E. & Fiorucci, S. The bile acid receptor FXR is a modulator of intestinal innate immunity. J. Immunol. 183, 6251–6261 (2009).

7. Pols, T. W. H. et al. Lithocholic acid controls adaptive immune responses by inhibition of Th1 activation through the Vitamin D receptor. PLOS ONE 12, e0176715 (2017).

8. Begley, M., Hill, C. & Gahan, C. G. M. Bile salt hydrolase activity in probiotics. Appl. Environ. Microbiol. 72, 1729–1738 (2006).

9. Chiang, J. Y. Recent advances in understanding bile acid homeostasis. F1000Res 6, 2029 (2017).

10. Song, Z. et al. Taxonomic profiling and populational patterns of bacterial bile salt hydrolase (BSH) genes based on worldwide human gut microbiome. Microbiome 7, 9 (2019).

11. Strelow, J. M. A Perspective on the Kinetics of Covalent and Irreversible Inhibition. SLAS Discov 22, 3–20 (2017).

12. Hamilton, J. P. et al. Human cecal bile acids: concentration and spectrum. Am. J. Physiol. Gastrointest. Liver Physiol. 293, G256–G263 (2007).

13. Roberts, A. B. et al. Development of a gut microbe-targeted nonlethal therapeutic to inhibit thrombosis potential. Nat. Med. 24, 1407–1417 (2018).

14. Rossocha, M., Schultz-Heienbrok, R., Moeller, von, H., Coleman, J. P. & Saenger, W. Conjugated bile acid hydrolase is a tetrameric N-terminal thiol hydrolase with specific recognition of its cholyl but not of its tauryl product. Biochem. 44, 5739–5748 (2005).

15. Huijghebaert, S. M. & Hofmann, A. F. Influence of the amino acid moiety on deconjugation of bile acid amidates by cholylglycine hydrolase or human fecal cultures. J. Lipid Res. 27, 742–752 (1986).

16. Yao, L. et al. A selective gut bacterial bile salt hydrolase alters host metabolism. eLife 7, 675 (2018).

17. Liu, Q. et al. Developing Irreversible Inhibitors of the Protein Kinase Cysteinome. Chemistry & Biology 20, 146–159 (2013).

18. Gehringer, M. & Laufer, S. A. Emerging and Re-Emerging Warheads for Targeted Covalent Inhibitors: Applications in Medicinal Chemistry and Chemical Biology. Journal of Medicinal Chemistry acs.jmedchem.8b01153 (2019). doi:10.1021/acs.jmedchem.8b01153

19. Wilson, A. J., Kerns, J. K., Callahan, J. F. & Moody, C. J. Keap calm, and carry on covalently. Journal of Medicinal Chemistry 56, 7463–7476 (2013).

20. Serafimova, I. M. et al. Reversible targeting of noncatalytic cysteines with chemically tuned electrophiles. Nature Chemical Biology 8, 471–476 (2012).

21. Henise, J. C. & Taunton, J. Irreversible Nek2 kinase inhibitors with cellular activity. Journal of Medicinal Chemistry 54, 4133–4146 (2011).

22. Xie, T. et al. Pharmacological targeting of the pseudokinase Her3. Nature Chemical Biology 10, 1006–1012 (2014).

23. Quintás-Cardama, A., Kantarjian, H., Cortes, J. & Verstovsek, S. Janus kinase inhibitors for the treatment of myeloproliferative neoplasias and beyond. Nature Reviews Drug Discovery 10, 127–140 (2011).

24. Cohen, M. S., Zhang, C., Shokat, K. M. & Taunton, J. Structural bioinformatics-based design of selective, irreversible kinase inhibitors. Science 308, 1318–1321 (2005).

25. Yang, W. et al. MX1013, a dipeptide caspase inhibitor with potent in vivo antiapoptotic activity. Br. J. Pharmacol. 140, 402–412 (2003).

26. Angliker, H., Wikstrom, P., Rauber, P. & Shaw, E. The synthesis of lysylfluoromethanes and their properties as inhibitors of trypsin, plasmin and cathepsin B. Biochem. J. 241, 871–875 (1987).

27. Garland, M. et al. Covalent Modifiers of Botulinum Neurotoxin Counteract Toxin Persistence. ACS Chem. Biol. 14, 76–87 (2019).

28. Miller, R. M. & Taunton, J. Targeting protein kinases with selective and semipromiscuous covalent inhibitors. Meth. Enzymol. 548, 93–116 (2014).

29. Coleman, J. P. & Hudson, L. L. Cloning and characterization of a conjugated bile acid hydrolase gene from Clostridium perfringens. Appl. Environ. Microbiol. 61, 2514–2520 (1995).

30. Tanaka, H., Hashiba, H., Kok, J. & Mierau, I. Bile salt hydrolase of Bifidobacterium longum-biochemical and genetic characterization. Appl. Environ. Microbiol. 66, 2502–2512 (2000).

31. Wang, Z. et al. Identification and Characterization of a Bile Salt Hydrolase from Lactobacillus salivarius for Development of Novel Alternatives to Antibiotic Growth Promoters. Appl. Environ. Microbiol. 78, 8795–8802 (2012).

32. Stellwag, E. J. & Hylemon, P. B. Purification and characterization of bile salt hydrolase from Bacteroides fragilis subsp. fragilis. Biochimica et Biophysica Acta (BBA) - Enzymology 452, 165–176 (1976).

33. Smith, K., Zeng, X. & Lin, J. Discovery of bile salt hydrolase inhibitors using an efficient high-throughput screening system. PLOS ONE 9, e85344 (2014).

34. Sayin, S. I. et al. Gut microbiota regulates bile acid metabolism by reducing the levels of tauro-beta-muricholic acid, a naturally occurring FXR antagonist. Cell Metab. 17, 225–235 (2013).

35. Hofmann, A. F. The function of bile salts in fat absorption. The solvent properties of dilute micellar solutions of conjugated bile acids. Biochem. J. 89, 57–68 (1963).

36. Kraal, L., Abubucker, S., Kota, K., Fischbach, M. A. & Mitreva, M. The prevalence of species and strains in the human microbiome: a resource for experimental efforts. PLOS ONE 9, e97279 (2014).

37. Li, F. et al. Microbiome remodelling leads to inhibition of intestinal farnesoid X receptor signalling and decreased obesity. Nat Commun 4, 2384 (2013).

38. Wallace, B. D. et al. Alleviating cancer drug toxicity by inhibiting a bacterial enzyme. Science 330, 831–835 (2010).

39. Joyce, S. A. et al. Regulation of host weight gain and lipid metabolism by bacterial bile acid modification in the gut. Proc. Natl. Acad. Sci. U.S.A. 111, 7421–7426 (2014).

40. Ma, C. et al. Gut microbiome-mediated bile acid metabolism regulates liver cancer via NKT cells. Science 360, eaan5931 (2018).

41. Kabsch, W. XDS. Acta Crystallogr. D Biol. Crystallogr. 66, 125–132 (2010).

42. McCoy, A. J. et al. Phaser crystallographic software. J Appl Crystallogr 40, 658–674 (2007).

43. Emsley, P. & Cowtan, K. Coot: model-building tools for molecular graphics. Acta Crystallogr. D Biol. Crystallogr. 60, 2126–2132 (2004).

44. Afonine, P. V. et al. Towards automated crystallographic structure refinement with phenix.refine. Acta Crystallogr. D Biol. Crystallogr. 68, 352–367 (2012).

45. Chen, V. B. et al. MolProbity: all-atom structure validation for macromolecular crystallography. Acta Crystallogr. D Biol. Crystallogr. 66, 12–21 (2010).

46. Morin, A. et al. Collaboration gets the most out of software. eLife 2, e01456 (2013).

47. Zhang, Z. & Marshall, A. G. A universal algorithm for fast and automated charge state deconvolution of electrospray mass-to-charge ratio spectra. J. Am. Soc. Mass Spectrom. 9, 225–233 (1998).

48. Ficarro, S. B., Alexander, W. M. & Marto, J. A. mzStudio: A Dynamic Digital Canvas for User-Driven Interrogation of Mass Spectrometry Data. Proteomes 5, 20 (2017).

49. Xie, C. et al. An Intestinal Farnesoid X Receptor-Ceramide Signaling Axis Modulates Hepatic Gluconeogenesis in Mice. Diabetes 66, 613–626 (2017).

