## Supplementary Information for "Development of a Covalent Inhibitor of Gut Bacterial Bile Salt Hydrolases"

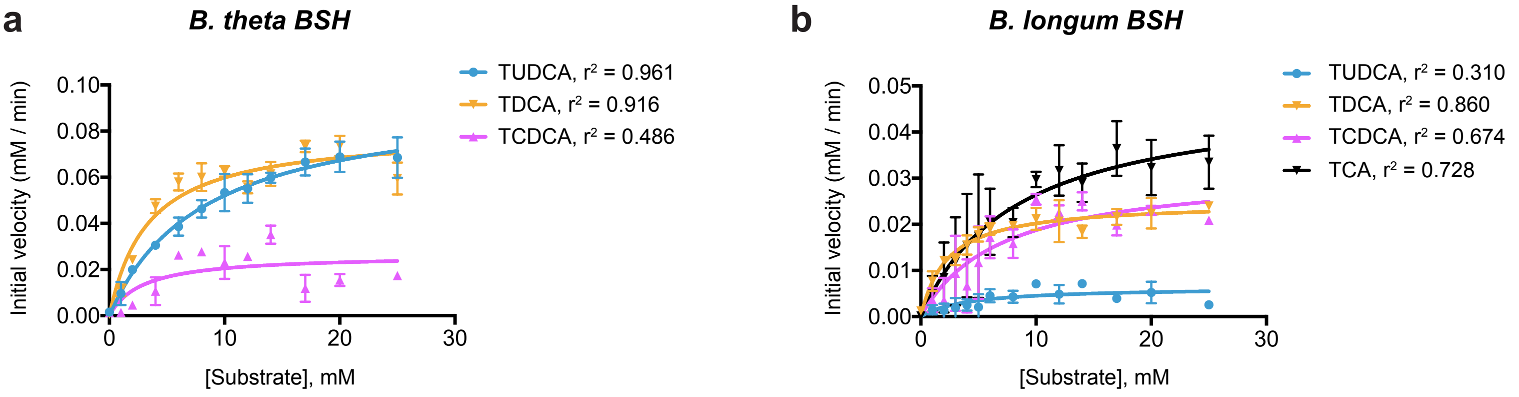

**Supplementary Figure 1.** Michaelis-Menten analysis of BSH kinetic data. Rate vs substrate concentration curves for *B. theta* BSH (**a**) and *B. longum* BSH (**b**). Each data point for both was obtained under initial velocity conditions in which less than 10% starting material was consumed. Assays were performed in biological triplicate. Graphpad was used to fit the Michaelis-Menten equations. Note: previously reported BSH kinetic parameters have been determined at the optimal pH of the enzymes (pH 4.2-6.5). We observed similar pH-dependent behavior for deconjugation by *B. theta* BSH and *B. longum* BSH. As a result, because all kinetic experiments herein were performed at physiological pH (7.5), lower kcat values were obtained for these two enzymes than typical literature values for similar enzymes.

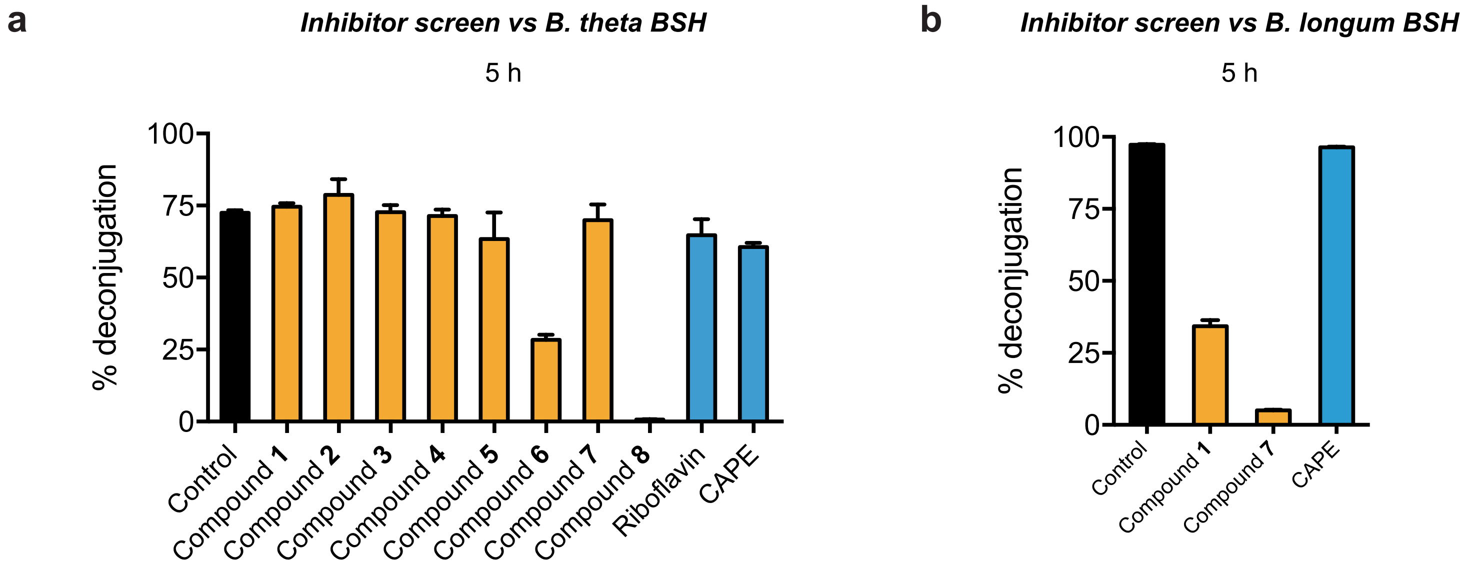

**Supplementary Figure 2.** Identification of compound **7** as a potent broad-spectrum BSH inhibitor. (**a**,**b**) Screen of inhibitors versus *B. theta* BSH (**a**) and B. longum BSH (**b**) showing % deconjugation at 5 hours. Inhibitor (100 μM) was incubated with 200 nM rBSH for 30 mins followed by addition of taurine-conjugated bile acid substrates (tauro-β-muricholic acid, TβMCA; tauro-cholic acid, TCA; tauro-ursodeoxycholic acid, TUDCA; and tauro-deoxycholic acid, TDCA, 25 μM each). Deconjugation of substrate was followed by UPLC-MS. Assays were performed in biological triplicate.

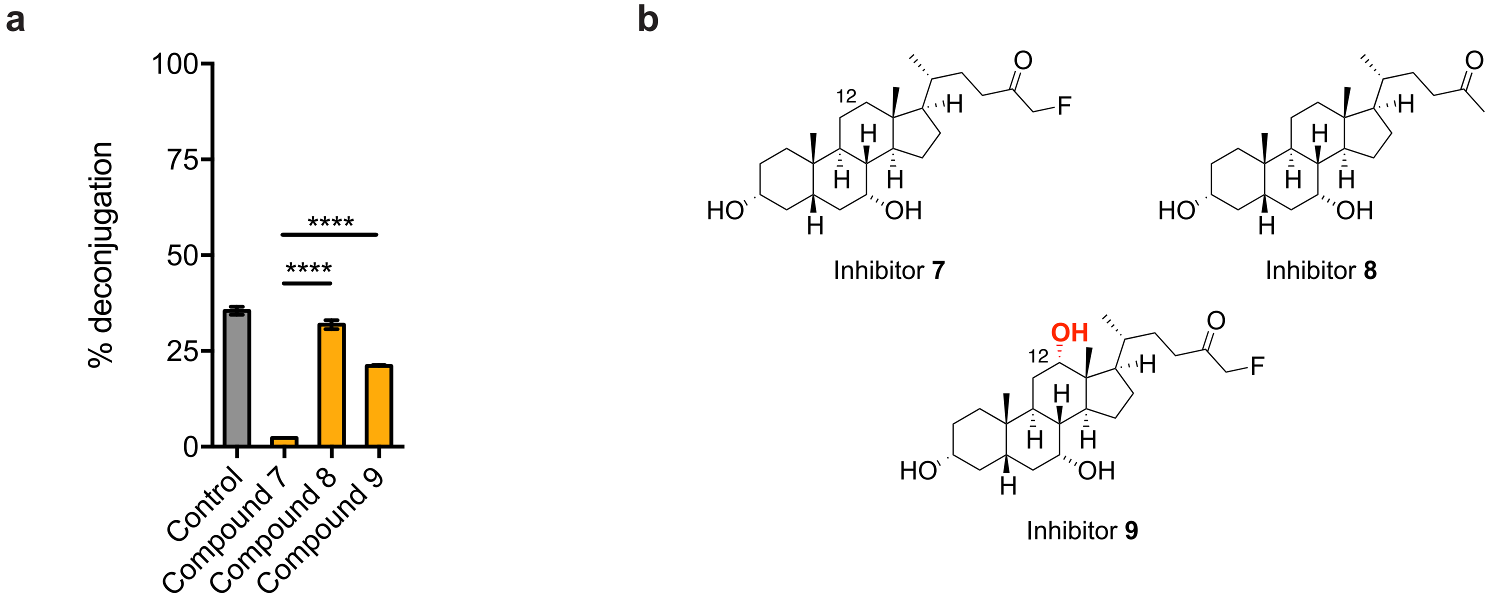

**Supplementary Figure 3. Compound structure affects BSH inhibitory activity against growing *B. theta* cultures.** (**a**) Compounds **8** and **9** are less potent inhibitors of *B. theta* BSH than compound **7**. Inhibitor (10 µM of compound **7**, **8**, or **9**) and taurine-conjugated bile acid substrates (TβMCA, TCA, TUDCA and TDCA, 25 µM each) were added to *B. theta* cultures at $OD_{600}$ 0.1. Cultures were allowed to grow into stationary phase and percent deconjugation at 24h was determined by UPLC-MS. One-way ANOVA followed by Tukey's multiple comparisons test. ****p<0.00001. Assays were performed in biological triplicate, and data are presented as mean ± SEM. (**b**) Structural comparison of compounds **7**, **8**, and **9**. Compound **8** lacks the α-FMK warhead, and compound **9** possesses a C12 = OH hydroxyl group.

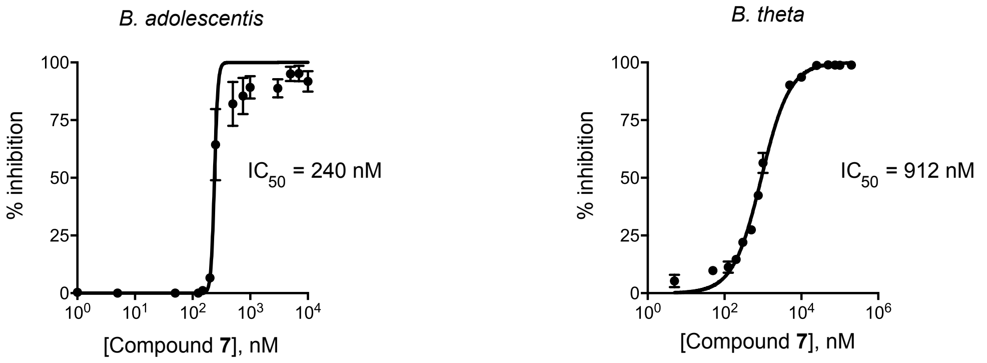

**Supplementary Figure 4. Compound 7 is a potent BSH inhibitor.** Dose-response curves and calculated $IC_{50}$ values for compound **7**. Pre-log phase cultures of *B. theta* (Gram negative) and *B. adolescentis* (Gram positive) were incubated with conjugated substrate (TUDCA or TDCA) and allowed to grow anaerobically for 48h and 24h, respectively. Deconjugation was quantified using UPLC-MS. Assays were performed in biological triplicate. Graphpad was used to fit $IC_{50}$ curves.

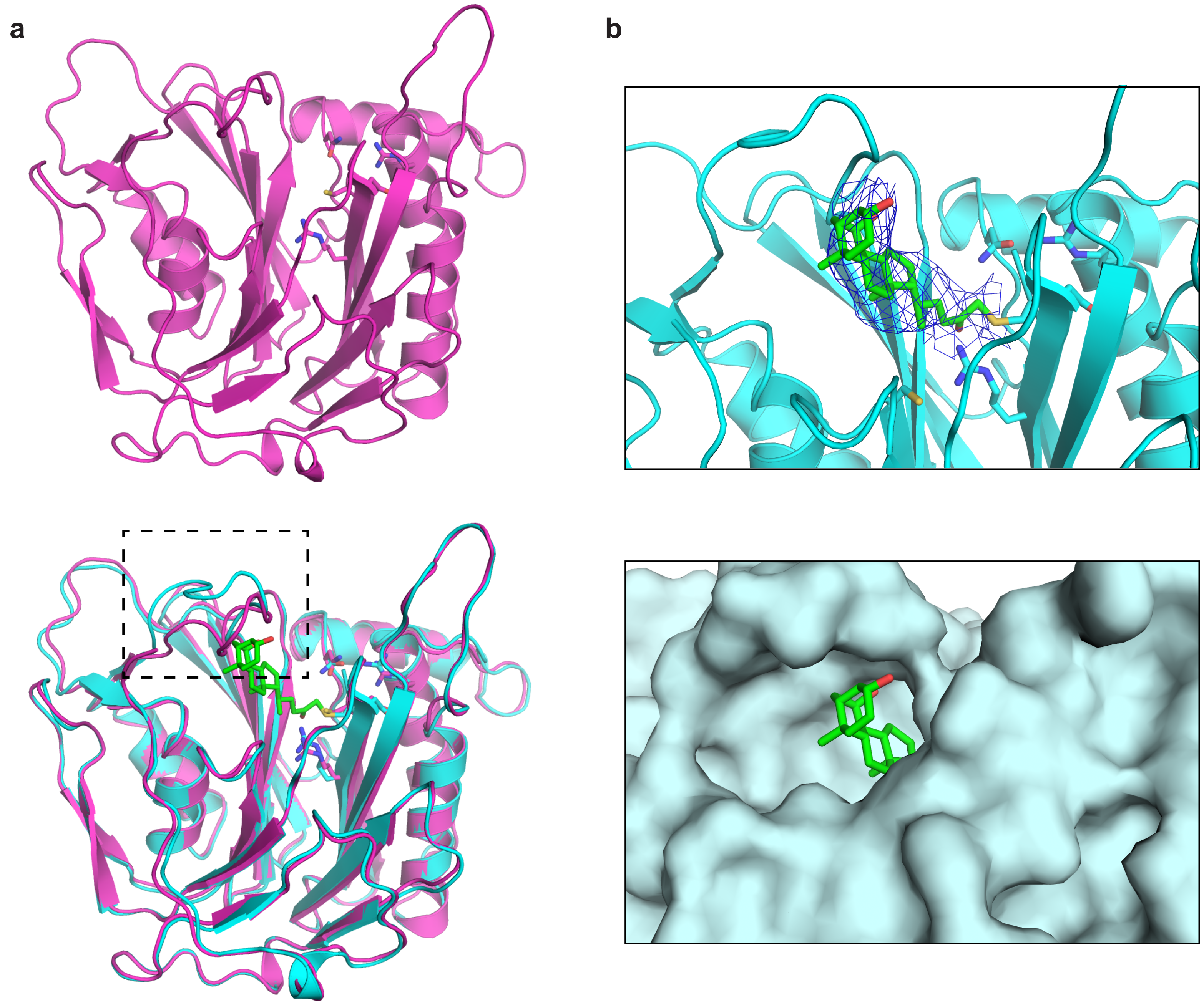

**Supplementary Figure 5.** Apo and co-crystal structures of *B. theta* BSH. (**a**) X-ray structure of *B. theta* BSH apoprotein (top) superimposed on the X-ray structure of *B. theta* BSH covalently bound to compound **7** (bottom). The BSHs (apo in magenta, co-crystal structure in cyan) are shown in ribbon representation, with indicated side chains (magenta or cyan, respectively, with heteroatoms in CPK colors) rendered as sticks. Compound **7** (green, with heteroatoms in CPK colors) is rendered in stick form. Box (dashed lines) indicates loop (residues 127-138) that has repositioned in the co-crystal structure. (**b**) Co-crystal structure of *B. theta* BSH and compound **7** shown in ribbon (top, with electron density of the compound shown as a blue net) and surface (bottom) representations. The A ring of compound **7** is solvent-exposed. Panels were prepared using PYMOL software (Schroedinger).

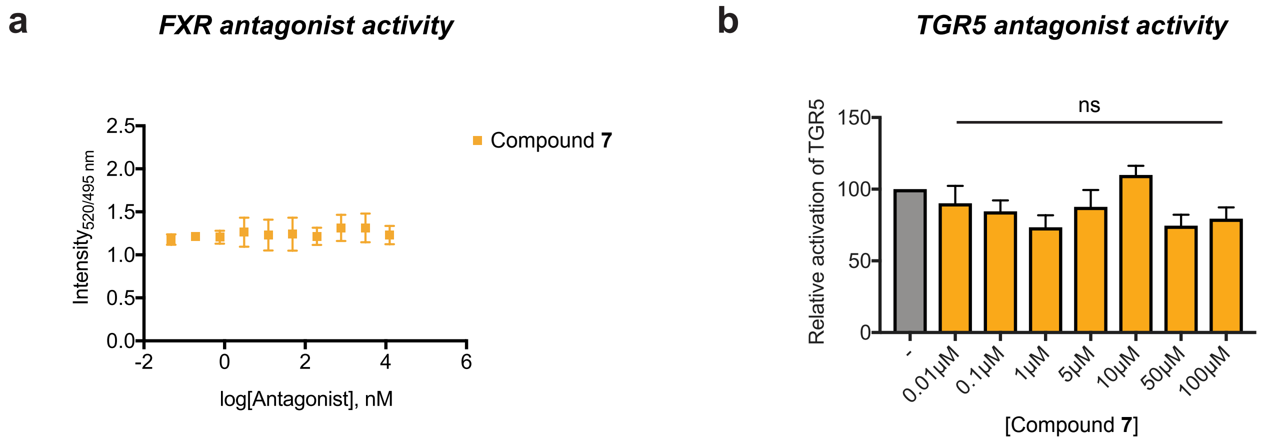

**Supplementary Figure 6.** Compound **7** is not an antagonist of FXR or TGR5. (**a**) Antagonist activity of compound **7** was evaluated in a coactivator recruitment assay in the presence of FXR agonist GW4064 at its EC50 value (50 nM, as determined in the corresponding agonist assay). n=4 biological replicates per concentration. (**b**) Endogenous TGR5 antagonist activity was measured by incubating Caco-2 cells with varying concentrations of compound **7** overnight in the presence of 10 μM of the TGR5 agonist LCA. n≥3 biological replicates per concentration. One-way ANOVA followed by Dunnett's multiple comparisons test, ns = not significant. All data are presented as mean ± SEM.

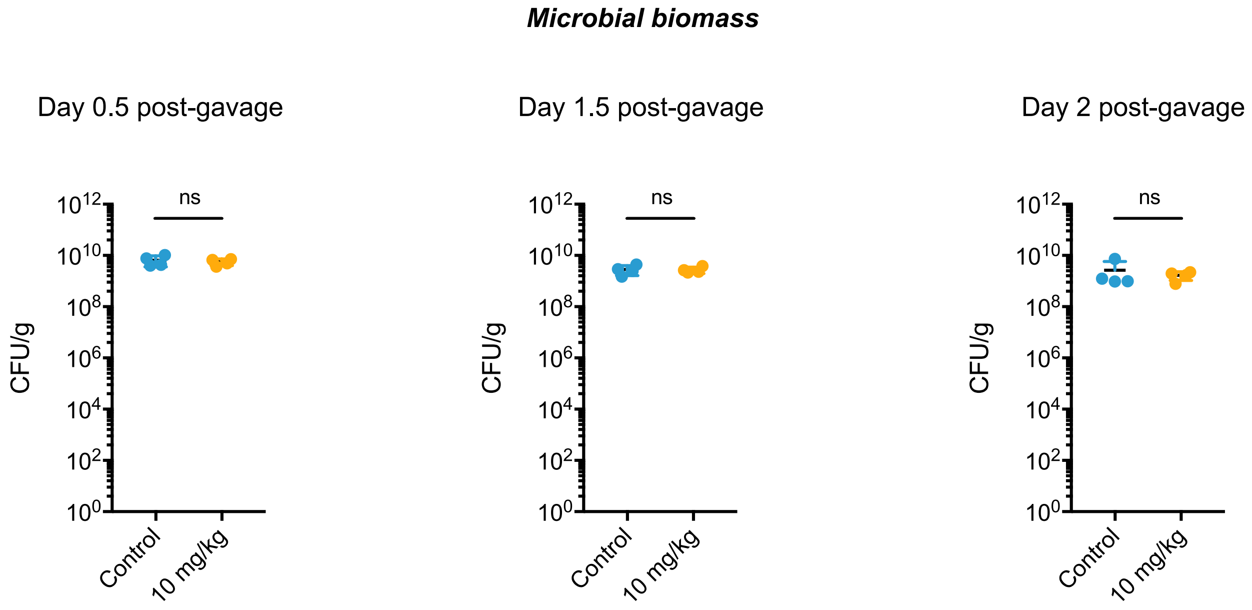

**Supplementary Figure 7.** Compound **7** did not affect microbial biomass in vivo. CFU/g did not differ between the inhibitor- and vehicle-treated groups 0.5, 1.5, or 2 days days post-gavage. n = 4 mice per group, Mann-Whitney test, ns = not significant.

**Supplementary Table 1. Crystallographic parameters.***

| Data Collection | BSH Compound | BSH |
|---|---|---|
| Wavelength (λ, Å) | 0.979 | 0.979 |
| Resolution range (Å) | 49.28 - 3.506 (3.528 - 3.406) | 46.16 – 2.706 (2.803 – 2.706) |
| Space group | $P2_12_12$ | $P2_12_12_1$ |
| Unit cell (Å, degrees) | 98.58, 99.52, 162.12, 90, 90, 90 | 84.88, 92.32, 194.25, 90, 90, 90 |
| Total reflections | 100568 (9882) | 177990 (17741) |
| Unique reflections | 20034 (1916) | 38728 (3864) |
| Multiplicity | 5.0 (5.2) | 4.6 (4.6) |
| Completeness (%) | 97.00 (95.32) | 91.08 (92.25) |
| Mean I/sigma(I) | 7.84 (0.86) | 4.28 (0.93) |
| Wilson B-factor | 142.4 | 38.21 |
| R-sym | 0.1394 (1.989) | 0.368 (2.097) |
| R-meas | 0.1561 (2.229) | 0.4163 (2.361) |
| $CC_{1/2}$ | 0.997 (0.59) | 0.967 (0.437) |
| Reflections used in refinement | 20032 (1914) | 38657 (3855) |
| Reflections used for R-free | 1840 (182) | 2012 (187) |
| R-work | 0.2438 (0.3606) | 0.2538 (0.3845) |
| R-free | 0.2925 (0.3767) | 0.2973 (0.4233) |
| Number of non-hydrogen atoms | 10246 | 10563 |
| Macromolecules | 10218 | 10319 |
| Solvent | 0 | 244 |
| Ligands | 28 | 0 |
| Protein residues | 1293 | 1307 |
| RMS (bonds, Å) | 0.004 | 0.002 |
| RMS (angles, degrees) | 0.45 | 0.52 |
| Ramachandran favored (%) | 89.23 | 96.92 |
| Ramachandran allowed (%) | 10.38 | 3.08 |
| Ramachandran outliers (%) | 0.39 | 0 |
| Rotamer outliers (%) | 0.53 | 1.05 |
| Clashscore | 51.61 | 8.81 |
| Average B-factor ($Å^2$) | 187.15 | 34.90 |
| Macromolecules | 187.24 | 35.01 |
| Solvent | 0 | 30.19 |
| Ligands | 153.63 | 0 |
| TLS | 23 | NA |

*Highest shell statistics are reported in parentheses.

**Supplementary Table 2. Primers for BSH gene amplification.**

| Protein | Primer | Sequence |
| --- | --- | --- |
| *B. theta* BSH | Bt_BSH_F | ATA GCT AGC ATG TGT ACG CGG GCG GTT TAC |
| *B. theta* BSH | Bt_BSH_R | ATC GCT CGA GCA TGA CTG GCG TTT CAA AC |
| *B. longum* BSH | Bl_BSH_F | GAT TGG CTA GCA TGT GCA CCG GCG TTC GT |
| *B.longum* BSH | Bl_BSH_R | GGG CTC GAG ACG TGC CAC TGA GAT TAA TTC |

**Supplementary Information, Synthetic Protocols and Compound Characterization.**

**General:** All anhydrous reactions were run under a positive pressure of argon or nitrogen. Anhydrous methylene chloride (DCM) and tetrahydrofuran (THF) were purchased from Sigma Aldrich. Silica gel column chromatography was performed using 60 Å silica gel (230−400 mesh). NMR spectra recorded in $CDCl_3$ used residual chloroform or TMS as the internal reference.

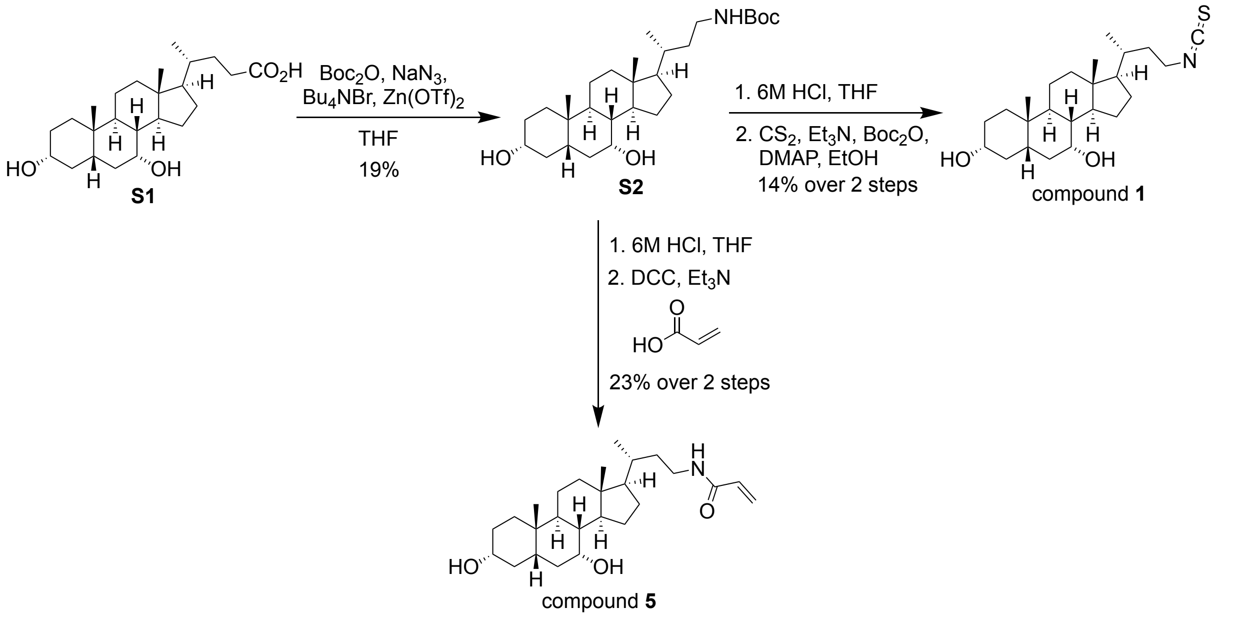

**Scheme 1. Synthesis of compounds 1 and 5.**

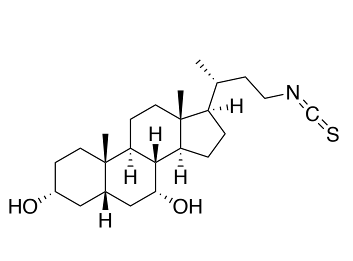

Compound **1.**

Step 1. To a solution of chenodeoxycholic acid (0.5 g, 1.27 mmol), sodium azide (0.29 g, 4.44 mmol), tetrabutylammonium bromide (61.0 mg, 0.19 mmol) and zinc trifluoromethanesulfonate (18.0 mg, 0.05 mmol) in 4.3 mL anhydrous THF at 40 °C was added di-tert-butyl dicarbonate (0.3 g, 1.399 mmol) and the mixture was heated overnight. The mixture was cooled to room temperature (rt) and quenched with 10 mL of 10% sodium nitrite and then diluted with 10 mL ethyl acetate. The organic layer was separated and the aqueous layer was extracted with ethyl acetate (2 x 10 mL). The combined organic layers where then dried over magnesium sulfate, filtered and concentrated on the rotovap. The crude compound was then purified by silica gel chromatography (80% ethyl acetate/20% hexanes) to provide compound **S2** (0.11 g, 19%) as a white foam.

Step 2. To the Boc amine **S2** (0.97 g, 2.09 mmol) in 4 mL THF, 1 mL of 6M HCl was added and the mixture was refluxed for 45 mins. The mixture was then cooled to rt and concentrated on the rotovap. The aqueous solution was then resuspended in 10 mL ethyl acetate and basified to pH 10 with 1M sodium hydroxide. The organic layer was separated and the aqueous layer was extracted with ethyl acetate (2 x 10 mL). The combined organic layers where then dried over sodium sulfate, filtered and concentrated to provide the free amine (0.51 g, 67%) which was used in the subsequent steps without further purification.

Step 3. A literature reported protocol was followed for the final step of the synthesis – *Tetrahedron Lett.,* **2008**, *49*, 3117-3119.

Briefly, to the amine (0.24 g, 0.66 mmol) in 2 mL ethanol, carbon disulphide (0.40 ml, 6.6 mmol) and trimethylamine (0.1 mL, 0.66 mmol) were added and the mixture was stirred at rt for 1h. The solution was then cooled to 0 °C and di-tert-butyl dicarbonate (0.14 g, 0.66 mmol) and DMAP (4.0 mg, 0.03 mmol) were added and the resulting mixture was stirred at 0 °C for 10 mins. The mixture was then warmed to rt and stirred for 10 mins following which it was concentrated on the rotovap. The crude compound was then purified by silica gel chromatography (75% ethyl acetate/25% hexanes) to provide the compound **1** (20.0 mg, 20%) as a clear oil.

**Compound 1.** TLC (Ethyl acetate:Hexanes, 85:15 v/v): $R_f$ = 0.5; $^1$H NMR (400 MHz, $CDCl_3$): δ 3.83 (s, 1H), 3.57-3.41 (m, 3H), 2.18 (q, *J* = 12.4 Hz, 1H), 1.99-1.79 (m, 7H), 1.71-1.64 (m, 3H), 1.57-1.11 (m, 14H), 1.00-0.81 (m, 7H), 0.67 (s, 3H); HRMS (*m/z*): $[M - 2H_2O + H]^+$ calcd. for $C_{24}H_{39}NO_2S$, 370.2568; found, 370.2543.

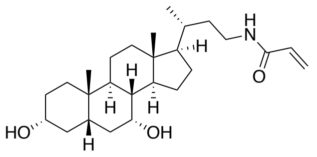

Compound **5.**

Step 1. The free amine was synthesized as per the conditions in step 2 for compound **1**.

Step 2. The acid (12.0 mg, 0.16 mmol) was dissolved in 1.65 mL anhydrous DCM followed by the addition of the coupling agent N,N'-dicyclohexylcarbodiimide (DCC) (42.0 mg, 0.20 mmol) and trimethylamine (76 µL, 0.55 mmol). The mixture was stirred at rt for 30 mins and then the free amine (50.0 mg, 0.14 mmol) dissolved in 1.4 mL DCM was added to the above mixture. The resulting solution was stirred at rt for 3 h. The mixture was then partitioned using 5 mL of 1M HCl and 5 mL DCM. The organic layer was separated and the aqueous layer was extracted with DCM (2 x 10 mL). The combined organic layers where then dried over sodium sulfate, filtered and concentrated. The crude compound was then purified by silica gel chromatography (90% ethyl acetate/10% hexanes) to provide the compound **5** (20.0 mg, 33%) as a clear oil.

**Compound 5.** TLC (Ethyl acetate:Hexanes, 80:20 v/v): $R_f$ = 0.12; $^1$H NMR (400 MHz, $CDCl_3$): δ 6.27 (dd, *J* = 16.8, 1.2 Hz, 1H), 6.07 (dd, *J* = 16.8, 10.4 Hz, 1H), 5.62 (dd, *J* = 10.4,

1.2 Hz, 1H), 5.45 (br s, 1H), 3.85-3.84 (m, 1H), 3.50-3.37 (m, 2H), 3.31-3.22 (m, 1H), 2.20 (q, *J* = 12.8 Hz, 1H), 2.01-1.79 (m, 5H), 1.73-1.10 (m, 19H), 1.01-0.94 (m, 4H), 0.90 (s, 3H), 0.66 (s, 3H); HRMS (*m/z*): $[M - 2H_2O + H]^+$ calcd. for $C_{26}H_{43}NO_3$, 382.3110; found, 382.3082.

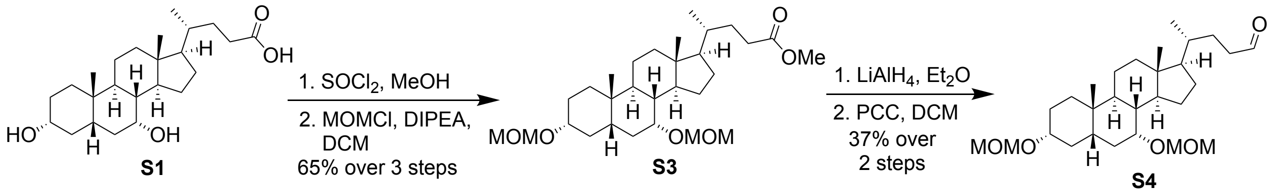

**Scheme 2. Synthesis of the common C-24 aldehyde intermediate.**

Step 1. To chenodeoxycholic acid (20.0 g, 50.8 mmol) suspended in 100 mL methanol at 0 °C, thionyl chloride (4.0 mL, 55.9 mmol) was added dropwise. The reaction was warmed to rt and stirred for 3 h. The reaction was quenched by the addition of 100 mL saturated sodium bicarbonate. The resulting mixture was then concentrated on the rotovap. The residue was partitioned between aqueous layer and 50 mL ethyl acetate. The organic layer was separated and the aqueous layer was extracted with ethyl acetate (2 x 50 mL). The combined organic layers where then dried over sodium sulfate, filtered and concentrated on the rotovap. The crude compound was then purified by silica gel chromatography (60% ethyl acetate/ 40% hexanes) to provide the methyl ester (20.6 g, quant.) as a white foam.

Step 2. The methyl ester (1.6 g, 3.93 mmol) was dissolved in 8 mL of anhydrous DCM and cooled to 0 °C under nitrogen. To this solution, *N,N*-diisopropylethylamine (1.4 mL, 11.79 mmol) was added followed by the slow addition of methoxymethyl chloride (1.2 mL, 11.79 mmol). The reaction mixture was then warmed to rt and stirred for 3 h. The reaction was quenched with the addition of 10 mL saturated sodium bicarbonate. The organic layer was separated and the aqueous layer was extracted with DCM (2 x 10 mL). The combined organic layers were then dried over sodium sulfate, filtered and concentrated. The crude compound was then purified by silica gel chromatography (25% ethyl acetate/75% hexanes) to provide the pure product **S3** (1.26 g, 65%) as a white foam.

Step 3. The protected methyl ester **S3** (1.73 g, 3.49 mmol) was dissolved in 14 mL anhydrous diethyl ether and cooled to 0 °C under nitrogen. LiAlH4 (0.27, 6.99 mmol) was added in portions to the above solution. The mixture was allowed to stir at 0 °C for 2 h and then quenched by the slow addition of 14mL of Rochelle's salt. The organic layer was separated and the aqueous layer was extracted with ethyl acetate (2 x 15 mL). The combined organic layers were then dried over magnesium sulfate, filtered and concentrated. The crude compound was then purified by silica gel chromatography (30% ethyl acetate/70% hexanes) to provide the pure C-24 alcohol (0.80 g, 50%) as a clear oil.

Step 4. To a suspension of pyridinium chlorochromate (1.42 g, 6.57 mmol) and silica gel (1.42 g) in 8 mL DCM at 0 °C, C-24 alcohol (1.9 g, 4.06 mmol) dissolved in another 8 mL DCM was added slowly. The resulting solution was then stirred at rt for 2 h. The reaction mixture was then filtered through a bed of celite and the residue was concentrated to provide the crude aldehyde **S4**. The crude compound was then purified by silica gel chromatography (20% ethyl acetate/80% hexanes) to provide pure aldehyde **S4** (1.39 g, 74%) as a clear oil.

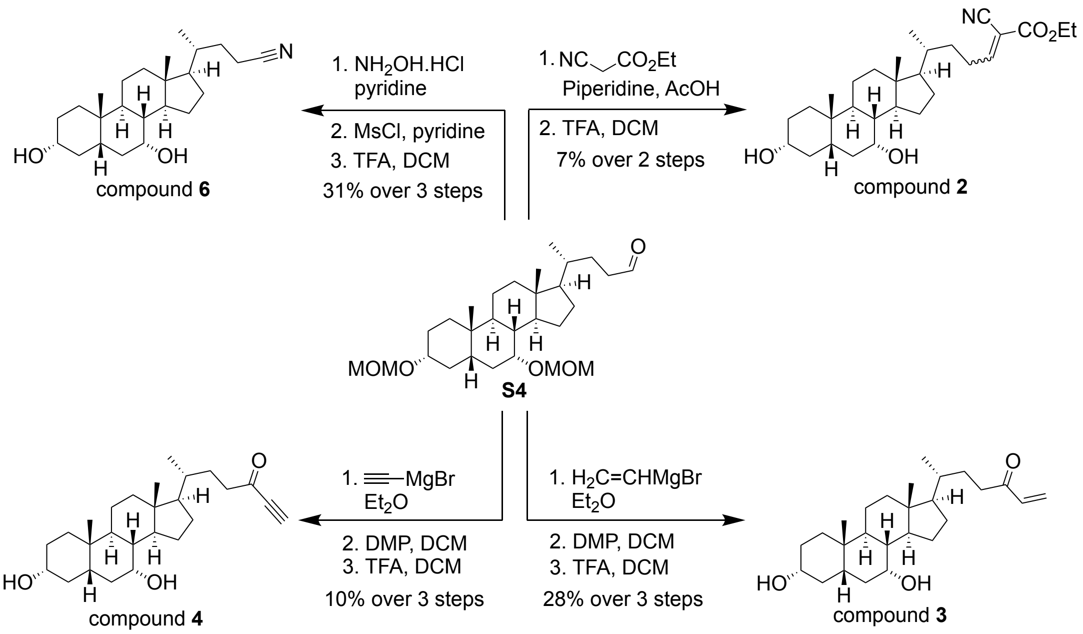

**Scheme 3. Synthesis of compounds 2-4 and 6.**

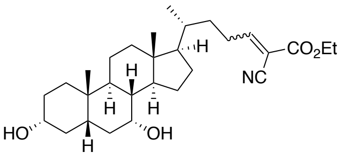

Compound **2.**

<u>Step 1.</u> A literature reported protocol was followed for this step of the synthesis – *J. Med. Chem.*, **2005**, *48*, 3026-3035.

To the aldehyde **S4** (0.12 g, 0.28 mmol) and ethyl 2-cyanoacrylate (35.0 mg, 0.30 mmol) at 0 °C, acetic acid (17 µL, 0.28 mmol) and piperidine (28 µL, 0.28 mmol) were added. The mixture was then stirred at rt overnight. The residue was then diluted with 5 mL DCM and washed with 5 mL of 1M HCl. The organic layer was separated and the aqueous layer was extracted with DCM (2 x 10 mL). The combined organic layers were then dried over sodium sulfate, filtered and concentrated. The crude compound was then purified by silica gel chromatography (30% ethyl acetate/70% hexanes) to provide the condensed intermediate (40.0 mg, 29%) as a clear oil.

Step 2. To the condensed product (50.0 mg, 0.09 mmol) in 1 mL DCM, 0.2 mL trifluoroacetic acid was added and the reaction mixture was stirred at 0 °C for 1 h. The mixture was then cooled to rt, diluted with 5 mL DCM and quenched slowly with 5 mL saturated sodium bicarbonate solution. The organic layer was separated and the aqueous layer was extracted with DCM (2 x 10 mL). The combined organic layers were then dried over sodium sulfate, filtered and concentrated. The crude compound was then purified by silica gel chromatography (60% ethyl acetate/40% hexanes) to provide the target compound **2** (10 mg, 24%) as a mixture of diastereomers which was used in the screen without further purification.

**Compound 2.** TLC (Ethyl acetate:Hexanes, 70:30 v/v): $R_f$ = 0.43; $^1$H NMR (400 MHz, $CDCl_3$): δ 7.64 (t, *J* = 8.0 Hz, 1H), 4.31 (q, *J* = 7.2 Hz, 2H), 3.85 (s, 1H), 3.50-3.44 (m, 1H), 2.63-2.43 (m, 2H), 2.20 (q, *J* = 12.8 Hz, 1H), 1.99-1.11 (m, 26H), 1.02-0.98 (m, 4H), 0.91 (s, 4H), 0.66 (s, 3H); HRMS (*m/z*): $[M + Na]^+$ calcd. for $C_{29}H_{45}NO_4$, 494.3246; found, 494.3233.

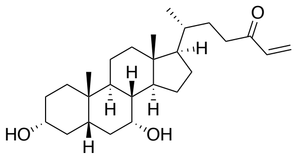

Compound **3.**

Step 1. In a flame dried flask, the aldehyde **S4** (0.50 g, 1.07 mmol) was dissolved in 4.3 mL diethyl ether under nitrogen. Vinylmagnesium bromide (1.6 mL, 1.60 mmol) was then added slowly to the above solution at 0 °C. The mixture was stirred at rt overnight. The reaction was quenched by the addition of 5 mL 1M HCl and diluted with 10 mL ethyl acetate. The organic layer was separated and the aqueous layer was extracted with ethyl acetate (2 x 10 mL). The combined organic layers were then dried over magnesium sulfate, filtered and concentrated.

The crude compound was then purified by silica gel chromatography (30% ethyl acetate/70% hexanes) to provide the alcohol as a mixture of diastereomers (0.40 g, 77%).

Step 2 and 3. Compound **3,** was synthesized following Steps 2 and 3 as described above for compound **4**. The overall yield is reported on the reaction scheme.

**Compound 3.** TLC (Ethyl acetate:Hexanes, 60:40 v/v): $R_f$ = 0.34; $^1$H NMR (400 MHz, $CDCl_3$): δ 6.35 (dd, *J* = 18.0, 10.8 Hz, 1H), 6.21 (dd, *J* = 17.6, 1.2 Hz, 1H), 5.89 (dd, *J* = 10.4, 0.8 Hz, 1H), 3.86-3.85 (m, 1H), 3.50-3.43 (m, 1H), 2.65-2.57 (m, 1H), 2.54-2.46 (m, 1H), 2.20 (q, *J* = 12.8 Hz, 1H), 2.01-1.11 (m, 22H), 1.02-0.91 (m, 9H), 0.66 (s, 3H); HRMS (*m/z*): $[M - 2H_2O + H]^+$ calcd. for $C_{26}H_{42}O_3$, 367.3001; found, 367.2985.

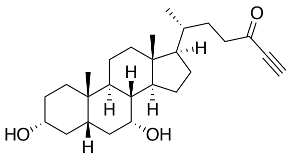

Compound **4**.

Step 1. In a flame dried flask, the aldehyde **S4** (0.20 g, 0.65 mmol) was dissolved in 1.5 mL diethyl ether under nitrogen. Ethynylmagnesium bromide (1.2 mL, 0.97 mmol) was then added slowly to the above solution at 0 °C. The mixture was stirred at rt overnight. The reaction was quenched by the addition of 2 mL 1M HCl and diluted with 10 mL ethyl acetate. The organic layer was separated and the aqueous layer was extracted with ethyl acetate (2 x 10 mL). The combined organic layers were then dried over magnesium sulfate, filtered and concentrated. The crude compound was then purified by silica gel chromatography (30% ethyl acetate/70% hexanes) to provide the alcohol as a mixture of diastereomers (0.12 g, 57%).

Step 2. To the alcohol (0.12 g, 0.24 mmol) in 2.5 mL anhydrous DCM at 0 °C, Dess-Martin periodinane (1.1 mL, 0.37 mmol) was added slowly. The reaction mixture was then stirred at rt until the reaction was complete by TLC. Upon consumption of the starting material, the reaction was quenched by the addition of 3 mL saturated sodium thiosulfate solution and 3 mL saturated sodium bicarbonate solution. The organic layer was separated and the aqueous layer was extracted with DCM (2 x 10 mL). The combined organic layers were then dried over sodium sulfate, filtered and concentrated. The crude compound was then purified by to provide the product (60.0 mg, 50%) as a clear oil.

Step 3. To the protected compound (20.0 mg, 0.04 mmol) in 1.0 mL DCM, trifluoroacetic acid (12 μL, 0.149 mmol) was added and the reaction mixture was stirred at 0 °C for 1 h. The mixture was then cooled to rt, diluted with 5 mL DCM and quenched slowly with 5 mL saturated sodium bicarbonate solution. The organic layer was separated and the aqueous layer was extracted with DCM (2 x 10 mL). The combined organic layers were then dried over sodium sulfate, filtered and concentrated. The crude compound was then purified by silica gel chromatography (50% ethyl acetate/50% hexanes) to provide the target compound **4** (5.5 mg, 34%) as a clear oil.

**Compound 4.** TLC (Ethyl acetate:Hexanes, 60:40 v/v): $R_f$ = 0.28; $^1$H NMR (400 MHz, $CDCl_3$): δ 3.86-3.85 (m, 1H), 3.50-3.43 (m, 1H), 3.20 (s, 1H), 2.66-2.48 (m, 2H), 2.20 (q, *J* = 12.8 Hz, 1H), 2.02-1.80 (m, 6H), 1.73-1.11 (m, 18H), 1.02-0.91 (m, 7H), 0.66 (s, 3H); HRMS (*m/z*): $[M - 2H_2O + H]^+$ calcd. for $C_{26}H_{40}O_3$, 365.2844; found, 365.2827.

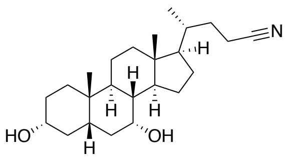

Compound **6**.

Step 1. To the aldehyde **S4** (0.50 g, 1.07 mmol) in 4 mL anhydrous pyridine, hydroxylamine hydrochloride (0.37 g, 5.37 mmol) was added and the reaction mixture was stirred at rt for 4 h. The mixture was then diluted with 20 mL DCM and washed with 20 mL 1M HCl. The organic layer was separated and the aqueous layer was extracted with DCM (2 x 20 mL). The combined organic layers were then dried over sodium sulfate, filtered and concentrated to provide the crude oxime which was used in the subsequent step without further purification.

Step 2. To the oxime (0.38 g, 0.79 mmol) in 4 mL pyridine, methanesulfonyl chloride (92 μL, 1.19 mmol) was added at 0 °C. The reaction mixture was warmed to rt and stirred for 18 h after which the mixture was diluted with 20 mL DCM and washed with 20 mL 1M HCl. The organic layer was separated and the aqueous layer was extracted with DCM (2 x 20 mL). The combined organic layers were then dried over sodium sulfate, filtered and concentrated to provide the crude oxime which was purified using silica gel chromatography (20% ethyl acetate/80% hexanes) to provide the pure bis-MOM protected nitrile (0.16g, 44% yield over two steps).

Step 3. To the protected nitrile (0.07 g, 0.151 mmol) in 1.5 mL tetrahydrofuran (THF), 50% HBr (100 μL, 0.607 mmol) was added and the reaction mixture was heated at 50 °C for 1 h. The mixture was then cooled to rt and diluted with 5 mL ethyl acetate and then quenched slowly with 5 mL saturated sodium bicarbonate solution. The organic layer was separated and the aqueous layer was extracted with ethyl acetate (2 x 20 mL). The combined organic layers were then dried over magnesium sulfate, filtered and concentrated. The crude compound was then purified by silica gel chromatography to provide the target compound **6** (40.0 mg, 71%) as a white solid.

**Compound 6.** TLC (Ethyl acetate:Hexanes, 60:40 v/v): $R_f$ = 0.2; $^1$H NMR (400 MHz, $CDCl_3$): δ 3.843-3.836 (m, 1H), 3.49-3.41 (m, 1H), 2.40-2.15 (m, 3H), 2.00-1.80 (m, 6H), 1.72-1.63 (m, 3H), 1.55-1.11 (m, 15H), 1.01-0.90 (m, 7H), 0.67 (s, 3H); HRMS (*m/z*): $[M - 2H_2O + H]^+$ calcd. for $C_{24}H_{39}NO_2$, 338.2848; found, 338.2828.

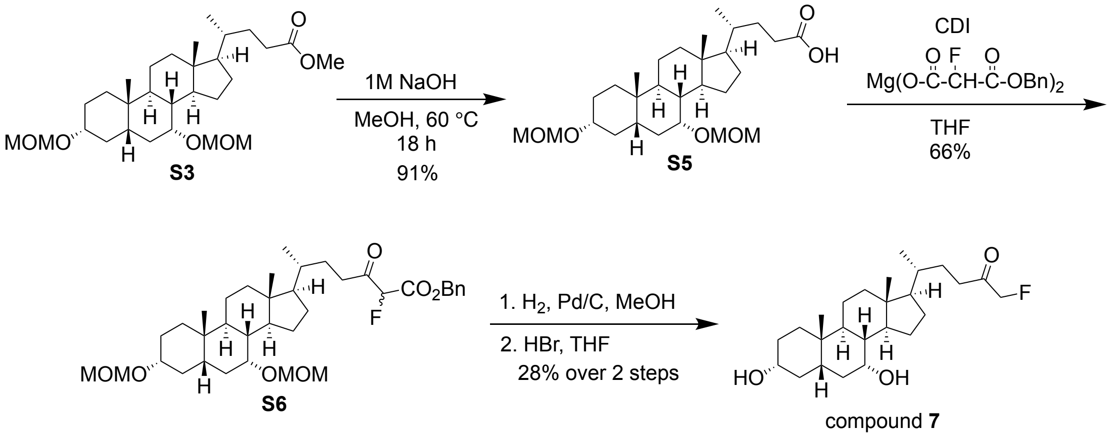

**Scheme 5. Synthesis of compound 7.**

The magnesium benzyl fluoromalonate coupling reagent was synthesized according to a reported protocol: James T Palmer. Process for Forming a Fluoromethyl Ketone. 5,210,272, May 11, 1993.

<u>Step 1.</u> To the protected methyl ester **S3** (6.4 g, 12.95 mmol) in 26 mL methanol, 28 mL 1M NaOH was added and the resulting solution was heated to 60 °C overnight. The mixture was then concentrated on the rotovap and resuspended in 30 mL each of 1M HCl and DCM. The organic layer was separated and the aqueous layer was extracted with DCM (2 x 30 mL). The combined organic layers were then dried over sodium sulfate, filtered and concentrated to provide the acid **S5** (5.7 g, 91%) as a white foam, which was used in subsequent reactions without further purification.

Step 2. To the C-24 acid **S5** (1.60 g, 3.33 mmol) in 6.5 mL of anhydrous THF, 1'-carbonyldiimidazole (CDI) (0.7 g, 4.33 mmol) was added and stirred at rt for 1 h. The magnesium benzyl fluoromalonate (1.20 g, 2.68 mmol) was suspended in 6.5 mL anhydrous THF and the above solution was added dropwise and the resulting mixture was stirred at rt for 18 h. The reaction was quenched by the addition of 10 mL of 1M HCl and concentrated on the rotovap. The residue was then partitioned using 10 mL DCM and 10 mL water. The organic layer was separated and the aqueous layer was extracted with DCM (2 x 10 mL). The combined organic layers were then dried over sodium sulfate, filtered and concentrated. The crude compound was then purified by silica gel chromatography (20% ethyl acetate/80% hexanes) to provide the pure compound **S5** (1.36 g, 66%) as a white foam.

Step 3. The compound **S5** (1.0 g, 1.63 mmol) and palladium on carbon (8.6 mg, 0.08 mmol) were suspended in 82 mL methanol. The flask for vacuumed and replaced with a hydrogen balloon. The reaction mixture was stirred at rt for 3 h. The solution was then filtered through a celite bed and the filtrate was concentrated to provide the bis-MOM protected fluoromethyl ketone (0.74 g, 96%).

Step 4. To the bis-MOM fluoromethyl ketone compound (0.74 g, 1.56 mmol) dissolved in 15.6 mL THF, 48% HBr (1.10 mL, 6.24 mmol) was added and the resulting solution was heated at 50 °C for 1 h. The mixture was cooled to rt and then quenched by slow addition of 20 mL saturated sodium bicarbonate solution. The biphasic solution was then concentrated on the rotovap and the resulting residue was partitioned using 20 mL DCM and 20 mL water. The organic layer was separated and the aqueous layer was extracted with DCM (2 x 20 mL). The combined organic layers were then dried over sodium sulfate, filtered and concentrated. The crude compound was then purified by silica gel chromatography (40% ethyl acetate/60%

hexanes to 60% ethyl acetate/40% hexanes) to provide the pure compound **7** (0.18 g, 29%) as a white foam.

**Compound 7.** TLC (Ethyl acetate:Hexanes, 70:30 v/v): $R_f$ = 0.36; $^1$H NMR (400 MHz, $CDCl_3$): δ 4.79 (d, *J* = 48.0 Hz, 1H), 3.85 (s, 1H), 3.50-3.44 (m, 1H), 2.60-2.44 (m, 2H), 2.20 (q. *J* = 13.2 Hz, 1H), 2.00-1.11 (m, 26H), 1.02-0.88 (m, 6H), 0.66 (s, 3H); HRMS (*m/z*): [M – $2H_2O$ + H]$^+$ calcd. for $C_{25}H_{41}FO_3$, 373.2907; found, 373.2888.

.

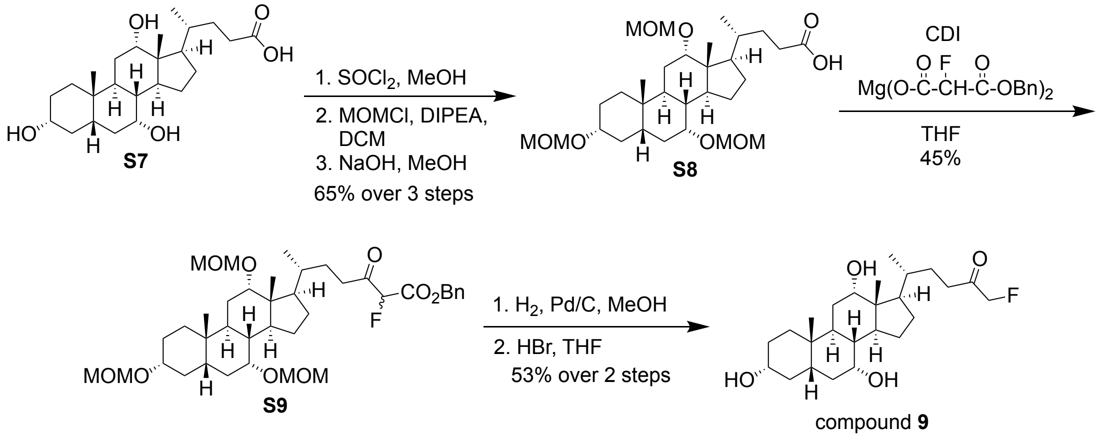

**Scheme 6. Synthesis of Compound 9.**

Compound **9** was synthesized from cholic acid (**S7**) as per the procedure described above for compound **7**. Yields for the synthesis of compound **9** are listed in the scheme above.

**Compound 9.** TLC (100% Ethyl acetate): $R_f$ = 0.12; $^1$H NMR (400 MHz, $CDCl_3$): δ 4.80 (d, *J* = 47.6 Hz, 1H), 3.96 (s, 1H), 3.85 (s, 1H),3.48-3.41 (m, 1H), 2.63-2.43 (m, 2H), 2.27-1.25 (m, 24H), 1.17-1.07 (m, 1H), 1.02-0.94 (m, 4H), 0.89 (s, 3H), 0.68 (s, 3H); HRMS (*m/z*): [M – $2H_2O$ + H]$^+$ calcd. for $C_{25}H_{41}FO_4$, 371.2750; found, 371.2725.

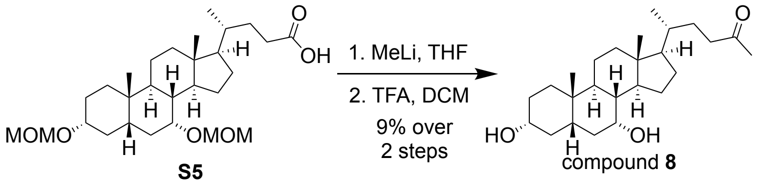

**Scheme 4. Synthesis of compound 8.**

Step 1. In a flame dried flask, the C-24 acid **S5** (0.36 g, 0.75 mmol) was dissolved in 7.5 mL anhydrous THF and cooled to -5 °C. Methyllithium (1.35 mL, 2.24 mmol) was then added dropwise and the reaction was stirred at -5 °C for 1 h. The reaction was quenched with 8 mL water and concentrated on the rotovap. The residue was then partitioned using ethyl acetate and water. The organic layer was separated and the aqueous layer was extracted with ethyl acetate (2 x 10 mL). The combined organic layers were then dried over sodium sulfate, filtered and concentrated. The crude compound was then purified by silica gel chromatography (30% ethyl acetate/70% hexanes) to provide the pure compound (0.15 g, 42%) as a white foam.

Step 2. Compound **8**, was synthesized following step 3 as described above for compound **4**. The overall yield is reported on the reaction scheme.

**Compound 8.** TLC (Ethyl acetate:Hexanes, 30:70 v/v): Rf = 0.1; $^1$H NMR (400 MHz, $CDCl_3$): δ 3.844-3.837 (m, 1H), 3.49-3.41 (m, 1H), 2.49-2.13 (m, 5H), 2.00-1.59 (m, 9H), 1.52-1.05 (m, 16H), 1.01-0.90 (m, 7H), 0.65 (s, 3H); HRMS (*m/z*): $[M - 2H_2O + H]^+$ calcd. for $C_{25}H_{42}O_3$, 355.3001; found, 355.2980.

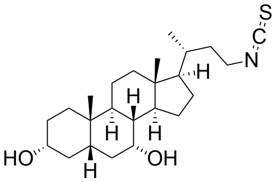

S
C
N
H
HO
OH

compound **1**

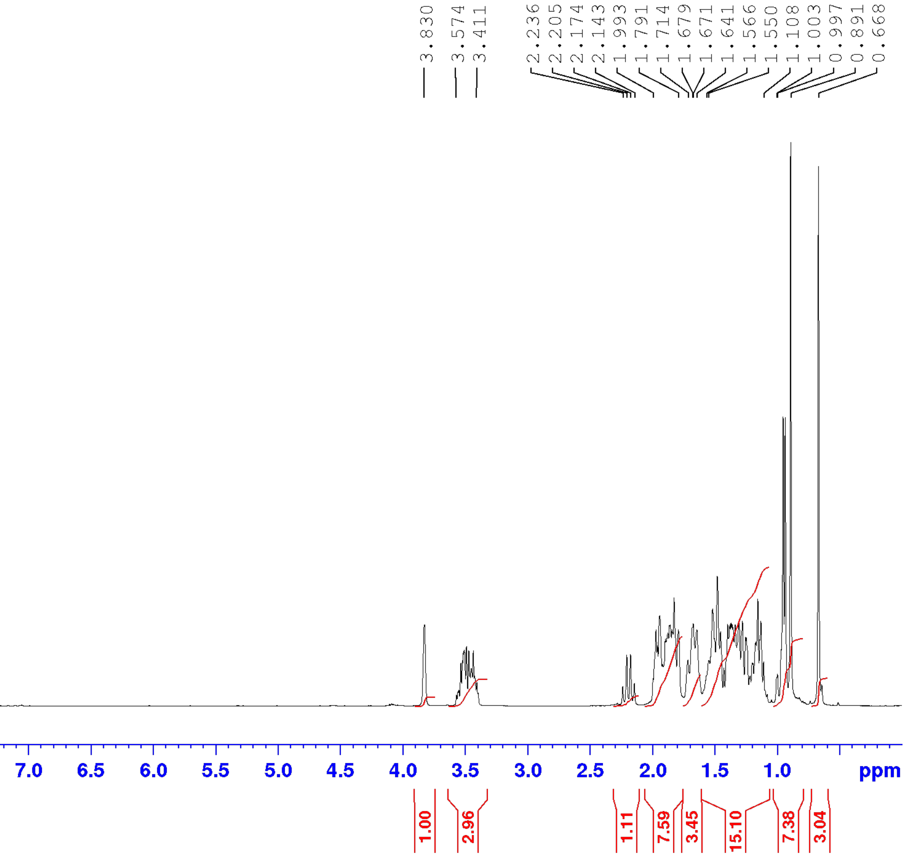

3.830
3.574
3.411
2.236
2.205
2.174
2.143
1.993
1.791
1.714
1.679
1.671
1.641
1.566
1.550
1.108
1.003
0.997
0.891
0.668
9.5
9.0
8.5
8.0
7.5
7.0
6.5
6.0
5.5
5.0
4.5
4.0
3.5
3.0
2.5
2.0
1.5
1.0
ppm
1.00
2.96
1.11
7.59
3.45
15.10
7.38
3.04

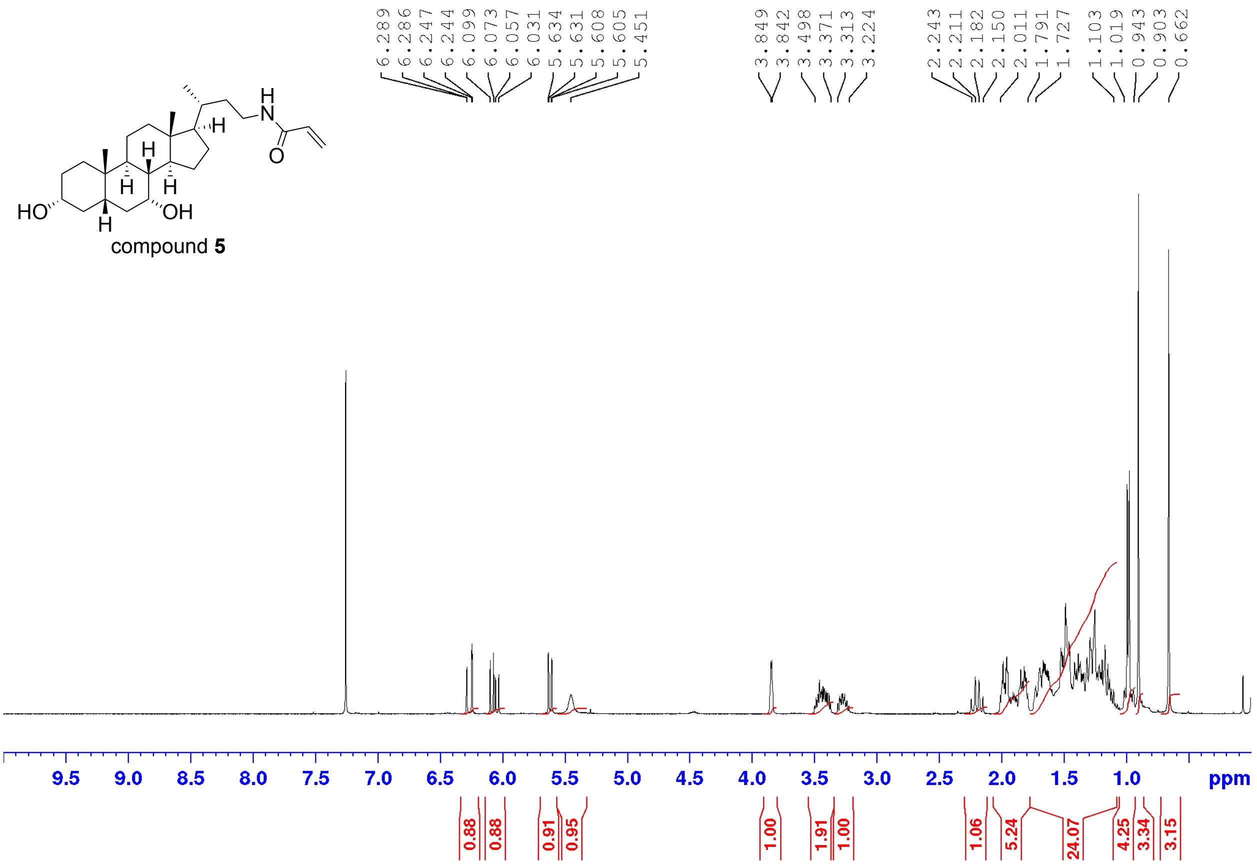
HO
OH
H
N
O
compound 5
6.289
6.286
6.247
6.244
6.099
6.073
6.057
6.031
5.634
5.631
5.608
5.605
5.451
3.849
3.842
3.498
3.371
3.313
3.224
2.243
2.211
2.182
2.150
2.011
1.791
1.727
1.103
1.019
0.943
0.903
0.662
9.5
9.0
8.5
8.0
7.5
7.0
6.5
6.0
5.5
5.0
4.5
4.0
3.5
3.0
2.5
2.0
1.5
1.0
ppm
0.88
0.88
0.91
0.95
1.00
1.91
1.00
1.06
5.24
24.07
4.25
3.34
3.15

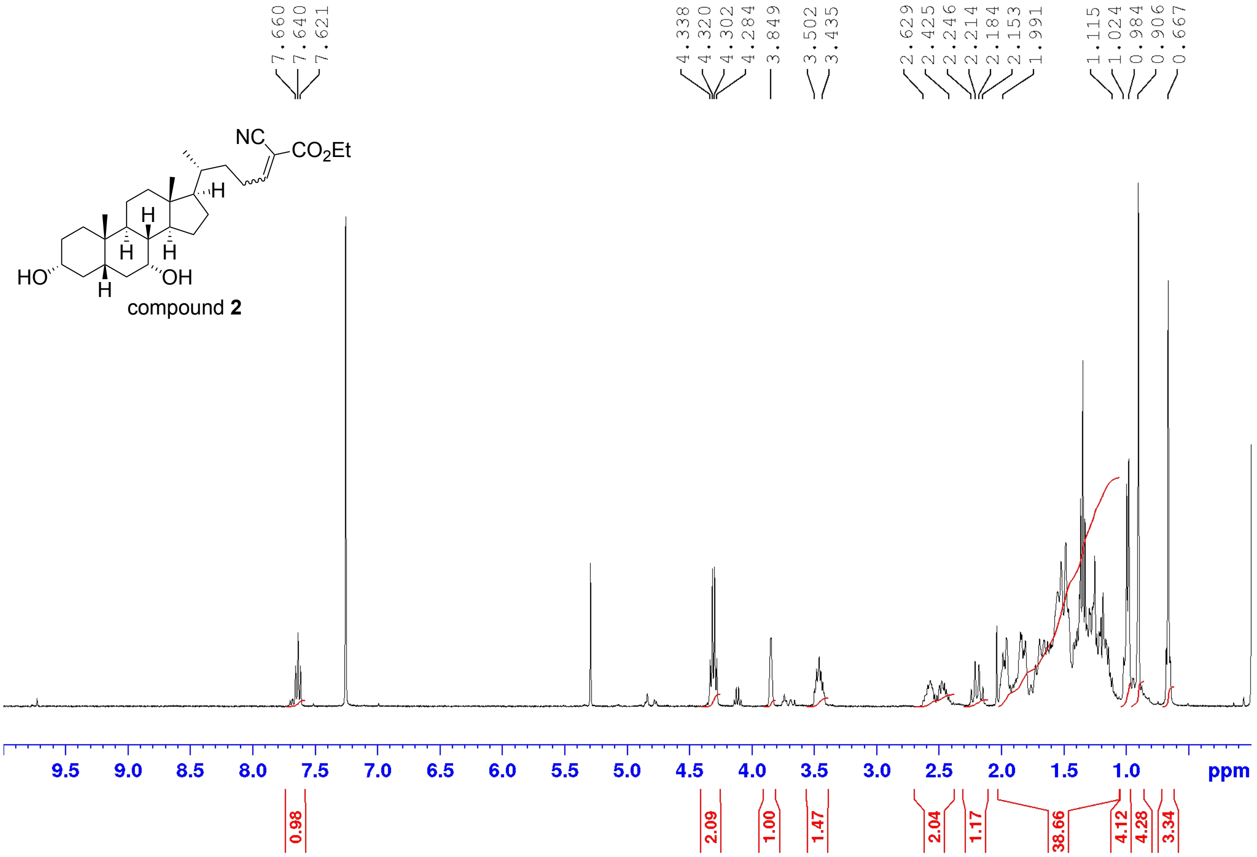
7.660
7.640
7.621
4.338
4.320
4.302
4.284
3.849
3.502
3.435
2.629
2.425
2.246
2.214
2.184
2.153
1.991
1.115
1.024
0.984
0.906
0.667
NC
$CO_2Et$
H
H
H
H
HO
OH
H
compound **2**
9.5
9.0
8.5
8.0
7.5
7.0
6.5
6.0
5.5
5.0
4.5
4.0
3.5
3.0
2.5
2.0
1.5
1.0
ppm
0.98
2.09
1.00
1.47
2.04
1.17
38.66
4.12
4.28
3.34

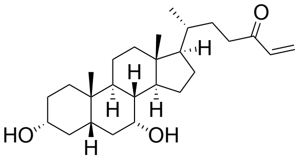
HO
OH
O
H
H
H
H
H

compound **3**

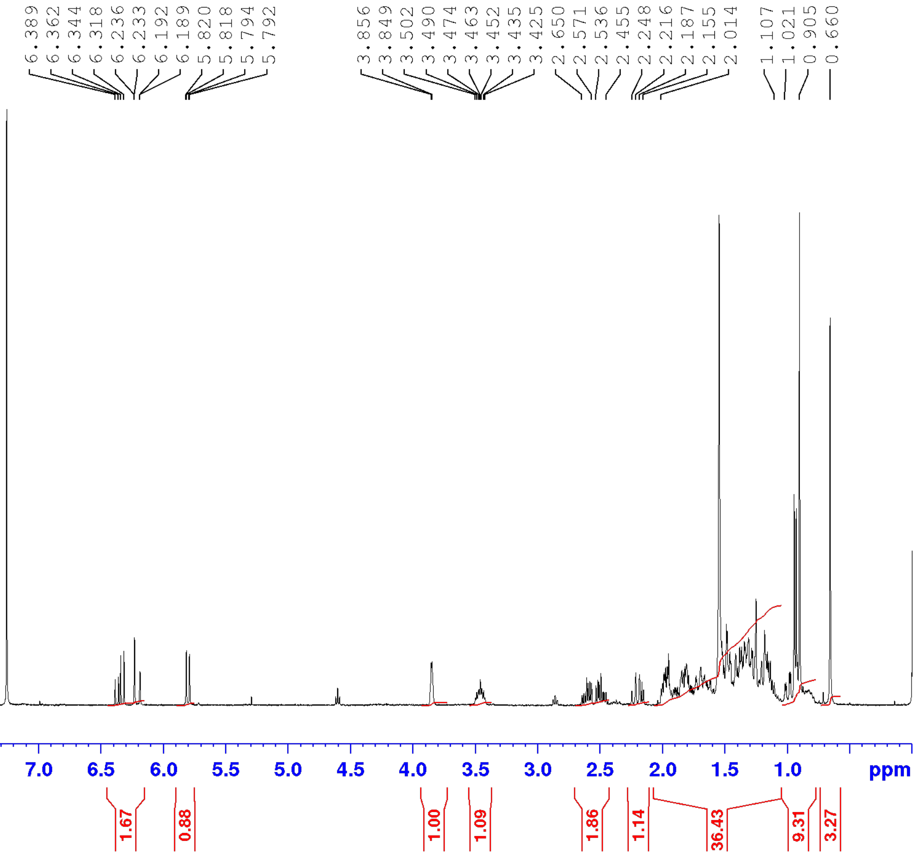
6.389
6.362
6.344
6.318
6.236
6.233
6.192
6.189
5.820
5.818
5.794
5.792
3.856
3.849
3.502
3.490
3.474
3.463
3.452
3.435
3.425
2.650
2.571
2.536
2.455
2.248
2.216
2.187
2.155
2.014
1.107
1.021
0.905
0.660
9.5
9.0
8.5
8.0
7.5
7.0
6.5
6.0
5.5
5.0
4.5
4.0
3.5
3.0
2.5
2.0
1.5
1.0
ppm
1.67
0.88
1.00
1.09
1.86
1.14
36.43
9.31
3.27

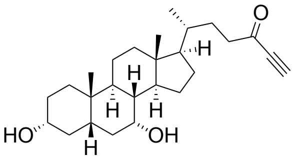

compound **4**

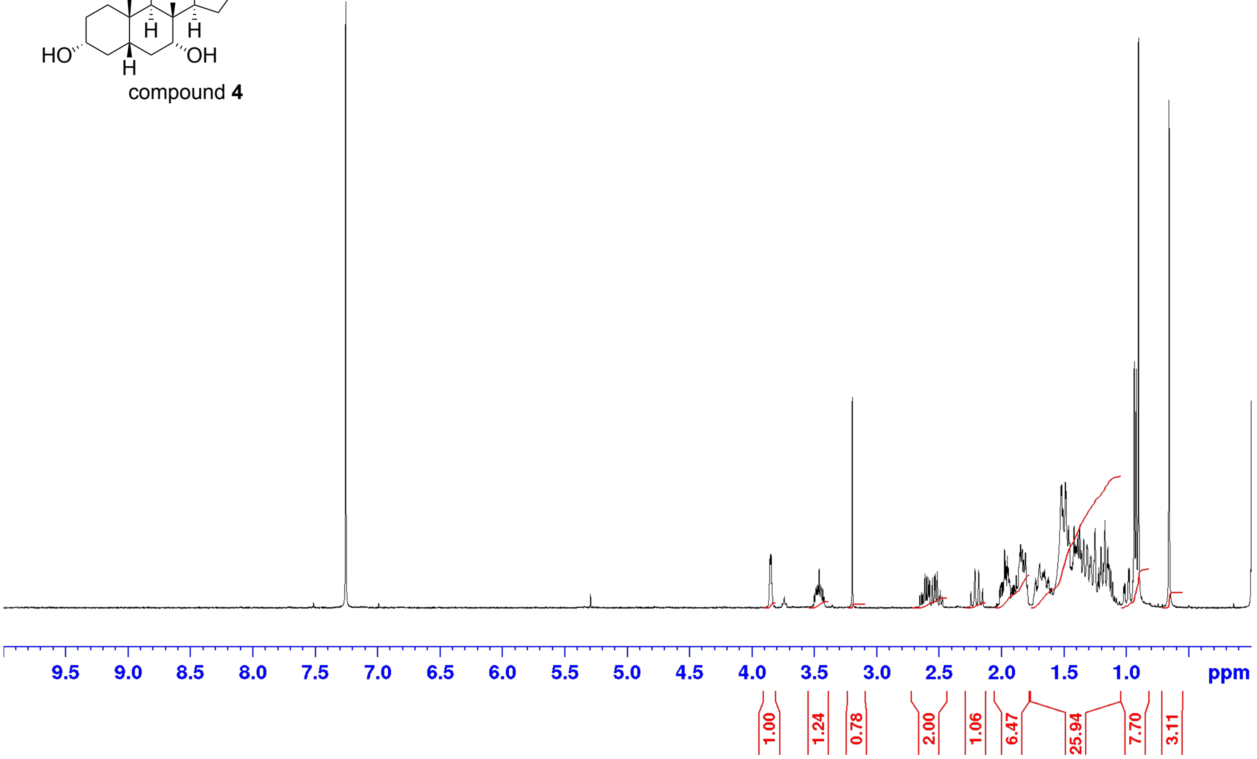

3.856
3.849
3.503
3.491
3.485
3.475
3.464
3.452
3.436
3.425
3.198
2.657
2.477
2.247
2.215
2.186
2.154
2.016
1.795
1.729
1.112
1.022
0.905
0.660
9.5 9.0 8.5 8.0 7.5 7.0 6.5 6.0 5.5 5.0 4.5 4.0 3.5 3.0 2.5 2.0 1.5 1.0 ppm
1.00
1.24
0.78
2.00
1.06
6.47
25.94
7.70
3.11

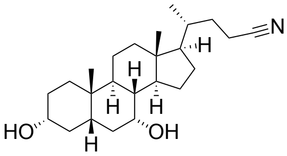

compound **6**

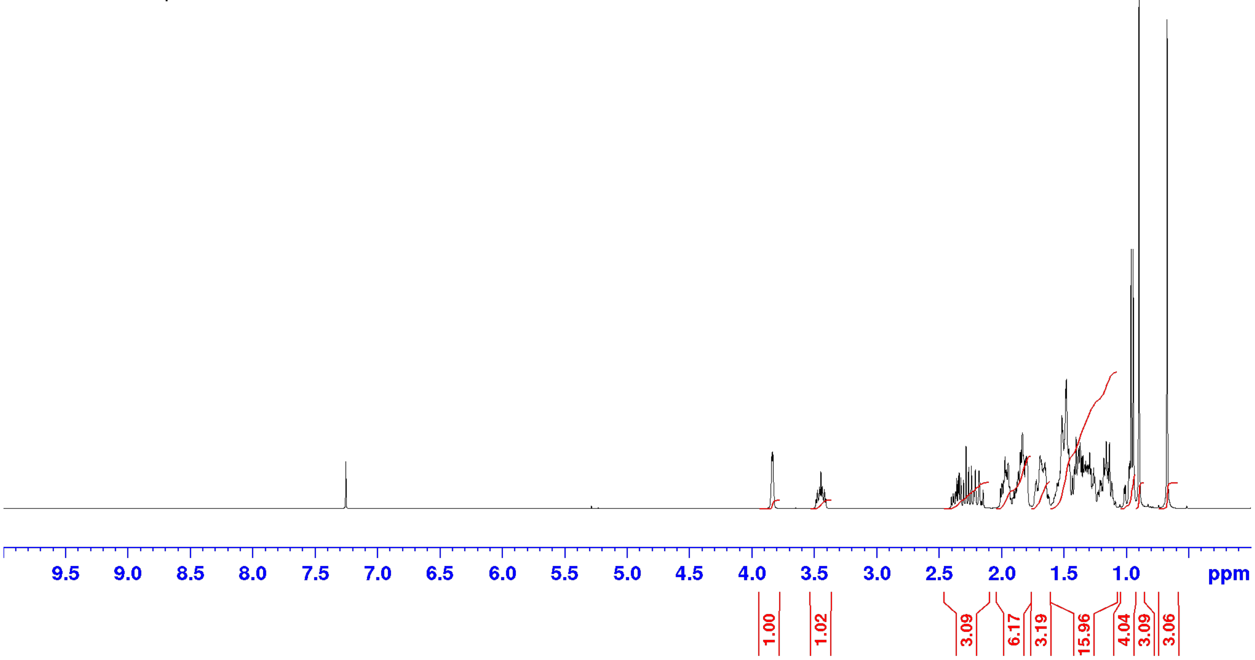

3.843
3.836
3.489
3.478
3.461
3.450
3.439
3.422
3.411
2.400
2.145
2.006
1.798
1.723
1.626
1.553
1.115
1.109
1.014
0.896
0.670
9.5
9.0
8.5
8.0
7.5
7.0
6.5
6.0
5.5
5.0
4.5
4.0
3.5
3.0
2.5
2.0
1.5
1.0
ppm
1.00
1.02
3.09
6.17
3.19
15.96
4.04
3.09
3.06

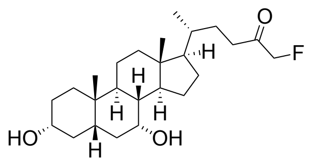
O
F
H
HO
OH

compound **7**

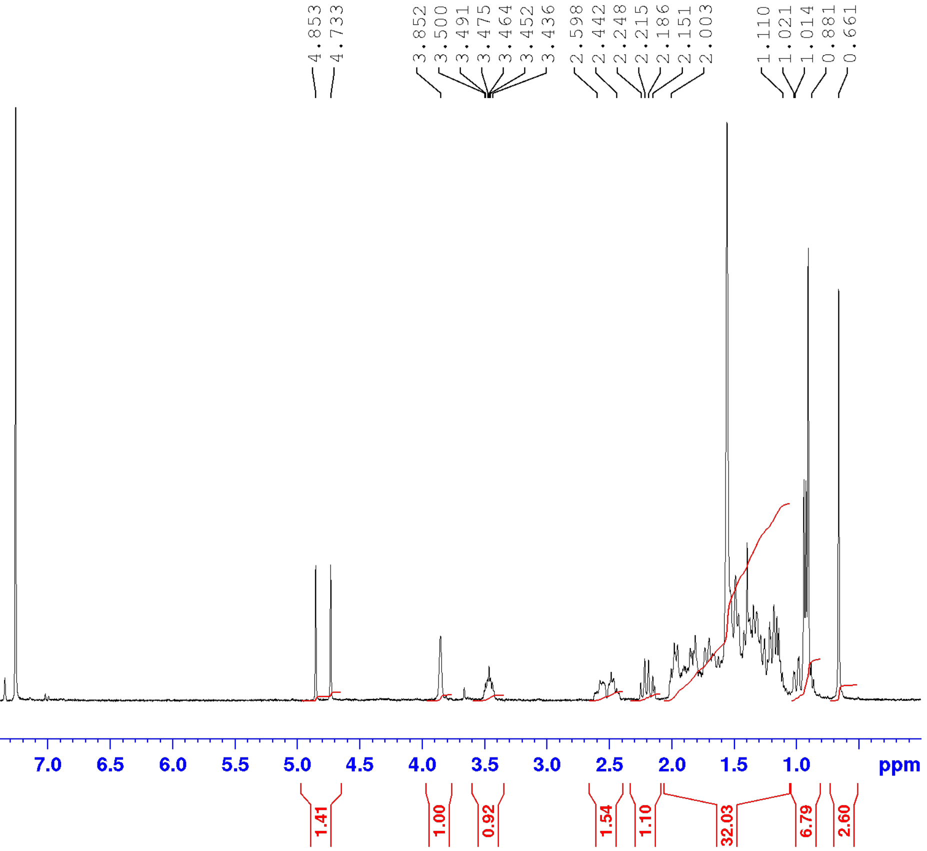
4.853
4.733
3.852
3.500
3.491
3.475
3.464
3.452
3.436
2.598
2.442
2.248
2.215
2.186
2.151
2.003
1.110
1.021
1.014
0.881
0.661
9.5 9.0 8.5 8.0 7.5 7.0 6.5 6.0 5.5 5.0 4.5 4.0 3.5 3.0 2.5 2.0 1.5 1.0 ppm
1.41
1.00
0.92
1.54
1.10
32.03
6.79
2.60

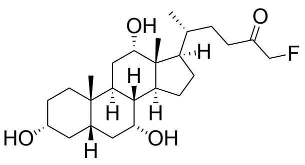

compound **9**

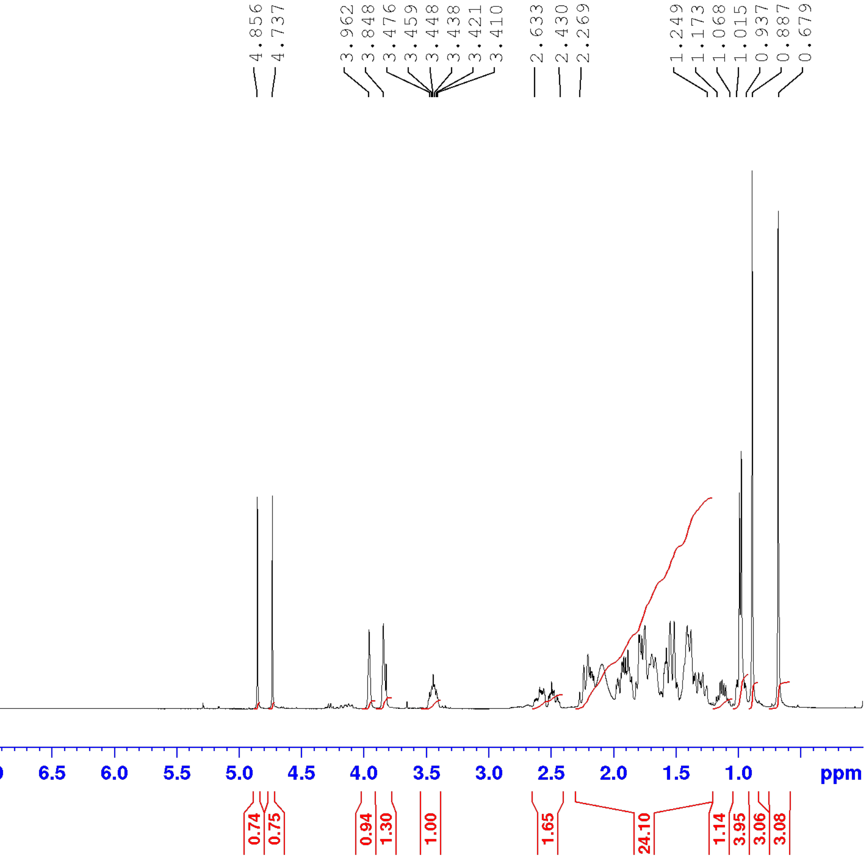

4.856
4.737
3.962
3.848
3.476
3.459
3.448
3.438
3.421
3.410
2.633
2.430
2.269
1.249
1.173
1.068
1.015
0.937
0.887
0.679
9.5
9.0
8.5
8.0
7.5
7.0
6.5
6.0
5.5
5.0
4.5
4.0
3.5
3.0
2.5
2.0
1.5
1.0
ppm
0.74
0.75
0.94
1.30
1.00
1.65
24.10
1.14
3.95
3.06
3.08

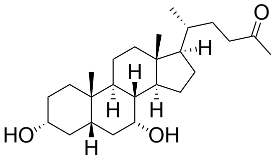

O
H
H
H
H
HO
H
OH

compound **8**

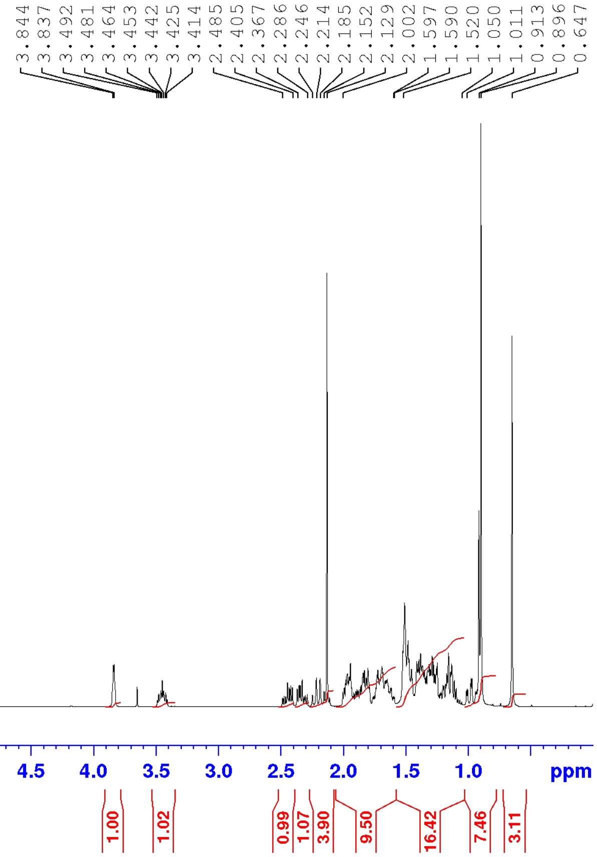

3.844
3.837
3.492
3.481
3.464
3.453
3.442
3.425
3.414
2.485
2.405
2.367
2.286
2.246
2.214
2.185
2.152
2.129
2.002
1.597
1.590
1.520
1.050
1.011
0.913
0.896
0.647
9.5
9.0
8.5
8.0
7.5
7.0
6.5
6.0
5.5
5.0
4.5
4.0
3.5
3.0
2.5
2.0
1.5
1.0
ppm
1.00
1.02
0.99
1.07
3.90
9.50
16.42
7.46
3.11
